# Chaperones directly and efficiently disperse stress-triggered biomolecular condensates

**DOI:** 10.1101/2021.05.13.444070

**Authors:** Haneul Yoo, Jared A.M. Bard, Evgeny Pilipenko, D. Allan Drummond

## Abstract

Heat shock triggers formation of intracellular protein aggregates and induction of a molecular disaggregation system. Although this system (Hsp100/Hsp70/Hsp40 in most cellular life) can disperse aggregates of model misfolded proteins, its activity on these model substrates is puzzlingly weak, and its endogenous heat-induced substrates have largely eluded biochemical study. Recent work has revealed that several cases of apparent heat-induced aggregation instead reflect evolved, adaptive biomolecular condensation. In budding yeast *Saccharomyces cerevisiae*, the resulting condensates depend on molecular chaperones for timely dispersal *in vivo*, hinting that condensates may be major endogenous substrates of the disaggregation system. Here, we show that the yeast disaggregation system disperses heat-induced biomolecular condensates of poly(A)-binding protein (Pab1) orders of magnitude more rapidly than aggregates of the most commonly used model substrate, firefly luciferase. Pab1 condensate dispersal also differs from aggregate dispersal in its molecular requirements, showing no dependence on small heat-shock proteins and a strict requirement for type II Hsp40. Unlike luciferase, Pab1 is not fully threaded (and thus not fully unfolded) by the disaggregase Hsp104 during dispersal, which we show can contribute to the extreme differences in dispersal efficiency. The Hsp70-related disaggregase Hsp110 shows some Pab1 dispersal activity, a potentially important link to animal systems, which lack cytosolic Hsp104. Finally, we show that the long-observed dependence of the disaggregation system on excess Hsp70 stems from the precise mechanism of the disaggregation system, which depends on the presence of multiple, closely spaced Hsp70s for Hsp104 recruitment and activation. Our results establish heat-induced biomolecular condensates of Pab1 as a direct endogenous substrate of the disaggregation machinery which differs markedly from previously studied foreign substrates, opening a crucial new window into the native mechanistic behavior and biological roles of this ancient system.

## Introduction

In all cellular life, a sudden increase in temperature—heat shock—causes formation of intracellular aggregates and production of heat shock proteins, many of which act as molecular chaperones (Parsell and Lindquist, 1993). An early and long-standing interpretation of these observations, which follow a wide range of so-called “proteotoxic stresses,” is that molecular chaperones are produced to protect cells from the toxic effects of stress-induced misfolded proteins and their aggregates (Lindquist, 1986; Morimoto, 2008; Vabulas et al., 2010).

Supporting this view, the molecular disaggregation system, which includes molecular chaperones Hsp100, Hsp70, and Hsp40, has been demonstrated to disperse aggregates of model substrates such as heat-misfolded firefly luciferase and restore their function *in vitro* (Glover and Lindquist, 1998; Goloubinoff et al., 1999). Decades of biochemical studies on these model substrates have uncovered the general mechanism of disaggregation as follows (reviewed in detail by Mogk et al. (2018)): First, J-domain proteins such as Hsp40 target Hsp70 to specific substrates while simultaneously stimulating Hsp70’s ATPase activity (Laufen et al., 1999; Lu and Cyr, 1998; Jiang et al., 2019; Faust et al., 2020). Second, substrate-bound Hsp70 recruits and de-represses the AAA+ disaggregase Hsp100 (Rosenzweig et al., 2013; Carroni et al., 2014; Seyffer et al., 2012; Haslberger et al., 2007). Third, Hsp100 threads substrate delivered by Hsp70 through its central channel to extract the substrate from aggregates (Haslberger et al., 2008; Gates et al., 2017; Avellaneda et al., 2020). Lastly, the threaded substrate is released from Hsp100 and undergoes either spontaneous or Hsp70/40-assisted folding to regain its native structure (Imamoglu et al., 2020).

A surprising and universal feature of biochemical studies of model misfolded substrate dispersal has been the use of—and in the case of Hsp70, a requirement for—substantial excesses of molecular chaperones over their substrates to achieve only limited dispersal. This is unlike a typical enzymatic reaction, although Hsp70 and Hsp104 are well-characterized enzymes and no chaperones are consumed during the disaggregation reaction.

An important possibility is that model substrates are not fully accurate models of endogenous substrates—and remarkably, the endogenous heat-induced substrates of the disaggregation system have largely eluded biochemical study. Alongside nascent polypeptides and prion fibers (Shorter and Lindquist, 2004; Inoue et al., 2004), heat-induced aggregates of misfolded mature proteins are considered major substrates of the disaggregation system. However, no endogenous mature protein has yet been identified to misfold in response to physiological heat shock in eukaryotes. As a consequence, a central element in our understanding of the heat shock response— that molecular chaperones directly engage and disperse endogenous aggregates induced by heat shock—has remained untested. A corollary is that the degree to which model thermolabile proteins, such as luciferase, accurately model endogenous aggregating proteins has been difficult to assess.

Recent work suggests that the proteotoxicity model, and the view that heat shock induces widespread protein misfolding, must be expanded. A proteome-wide study in budding yeast showed that a specific set of mature proteins form fully reversible aggregates in response to sublethal heat shock (Wallace et al., 2015). Closer inspection revealed that several cases of this apparent aggregation reflect evolved, adaptive biomolecular condensation (Riback et al., 2017; Iserman et al., 2020). For example, physiological heat shock temperature and pH changes cause poly(A)-binding protein (Pab1), an abundant and broadly conserved eukaryotic RNA-binding protein, to phase separate and form gel-like condensates *in vitro* (Riback et al., 2017). Suppressing Pab1 condensation reduces cell fitness during prolonged heat stress, indicating that condensation is adaptive (Riback et al., 2017). Similarly, heat-induced phase separation of translation initiation factor and DEAD-box helicase Ded1 confers an adaptive benefit to cells by promoting translational switch from housekeeping to stress-induced transcripts (Iserman et al., 2020). As illustrated by these studies, heat-induced biomolecular condensates of endogenous, mature proteins appear to be fundamentally different from misfolded protein aggregates in both mechanism of formation and, most importantly, fitness consequences.

Here we will use the term biomolecular condensates to refer to endogenous membraneless structures of concentrated biomolecules (Banani et al., 2017) regardless of the condensation mechanism, reserving the term phase separation for cases where it has been shown. We use the term aggregates to refer to amorphous clumps of misfolded proteins, which are commonly deleterious to cells,(Geiler-Samerotte et al., 2011) and which differ from endogenous condensates whose fitness consequences are adaptive in several cases.

Substantial *in vivo* evidence indicates that endogenous heat-induced condensates interact with the disaggregation system. All members of the yeast disaggregation system (Hsp104/Hsp70/Hsp40) co-localize with stress granules, which contain both Pab1 and Ded1 (Cherkasov et al., 2013; Walters et al., 2015; Kroschwald et al., 2015, 2018). Deletion or inhibition of any member of the system, or the Hsp70 nucleotide exchange factor (NEF) Hsp110 (Sse1/2), delays dissolution of stress granules during stress recovery (Cherkasov et al., 2013; Walters et al., 2015; Kroschwald et al., 2015, 2018). Interestingly, dispersal of endogenous stress granules precedes dispersal of exogenously expressed misfolded protein aggregates (Cherkasov et al., 2013; Kroschwald et al., 2015) and only the former correlates with the resumption of translation activity and the cell cycle (Cherkasov et al., 2013; Kroschwald et al., 2018).

We and others have hypothesized that heat-induced biomolecular condensates are major endogenous substrates of molecular chaperones (Wallace et al., 2015; Riback et al., 2017; Kroschwald et al., 2018; Yoo et al., 2019; Triandafillou et al., 2020; Begovich and Wilhelm, 2020; Snead and Gladfelter, 2019). However, the questions of whether molecular chaperones directly engage heat-induced biomolecular condensates, and whether and how functional engagement differs between adaptive condensates and aggregates of model misfolded substrates, have remained unanswered.

Here, we address these major open questions by reconstituting *in vitro* the dispersal of heat-induced Pab1 condensates by their cognate disaggregation system. We use independent methods to demonstrate that Hsp104, Hsp70, and the type II Hsp40 Sis1 are necessary and sufficient for complete dispersal of Pab1 condensates back to functional monomers. Comparative studies of Pab1 condensates and aggregates of misfolded luciferase reveal four key differences. First, and most strikingly, chaperones which show slow and incomplete dispersal of luciferase aggregates disperse Pab1 condensates rapidly and completely. Second, unlike luciferase (Cashikar et al., 2005), Pab1 does not require co-condensation with small heat shock protein Hsp26 for subsequent efficient dispersal. Third, unlike luciferase for which type I (Ydj1) and type II (Sis1) Hsp40 show synergistic activity (Nillegoda et al., 2015, 2017), Pab1 condensate dispersal depends only on Sis1 and is antagonized by Ydj1. Fourth, we show that unlike luciferase, Pab1 is only partially threaded by Hsp104 and readily regains its function upon dispersal.

Finally, we investigate the dispersal system’s puzzling dependence on excess Hsp70 for optimal activity, which we find also applies to Pab1 condensate dispersal. Combining biochemical experiments with modeling, we show that the required presence of multiple, closely-spaced Hsp70s for Hsp104 recruitment and activation suffices to render the disaggregation system sensitive to the relative Hsp70 level.

Our results establish heat-induced biomolecular condensates of Pab1 as direct endogenous substrates of the disaggregation system, and reveal that many important conclusions drawn from studying aggregates of “model” misfolded proteins do not generalize to endogenous condensates. Whether the remarkable efficiency of Pab1 dispersal is itself a general feature of native substrates remains to be seen. Further study of how chaperones engage with adaptive, endogenous substrates, and how this engagement differs from foreign or proteotoxic substrates, appears likely to yield substantial new insights into the mechanistic features and biological roles of this ancient molecular system.

## Results

### Heat shock causes Pab1 condensation, which is not spontaneously reversible

In budding yeast, Pab1 forms RNase-resistant, sedimentable condensates after physiological heat shock (Wallace et al., 2015; Riback et al., 2017). Condensates of mature, endogenous proteins disperse to their pre-stress soluble form within an hour, without degradation, as cells recover at 30°C (Wallace et al., 2015; Cherkasov et al., 2013).

Consistent with previous results, 20 minutes of heat shock at 42°C caused a roughly five-fold increase in the proportion of large sedimentable Pab1 compared to the pre-shock level (Figure 1A-C). This fraction decreased as the cells recovered at 30°C and reached the pre-shock level by 60 minutes. RNase treatment to release sedimentable species formed by RNA-protein interaction decreased the fraction of Pab1 sedimented at 100,000 g spin, but did not change the fraction of Pab1 sedimented at 8,000 g spin (Figure 1B), as previously reported (Riback et al., 2017). These results confirm that under these conditions Pab1 rapidly forms RNase-resistant assemblies which persist upon return to pre-shock temperatures *in vivo*.

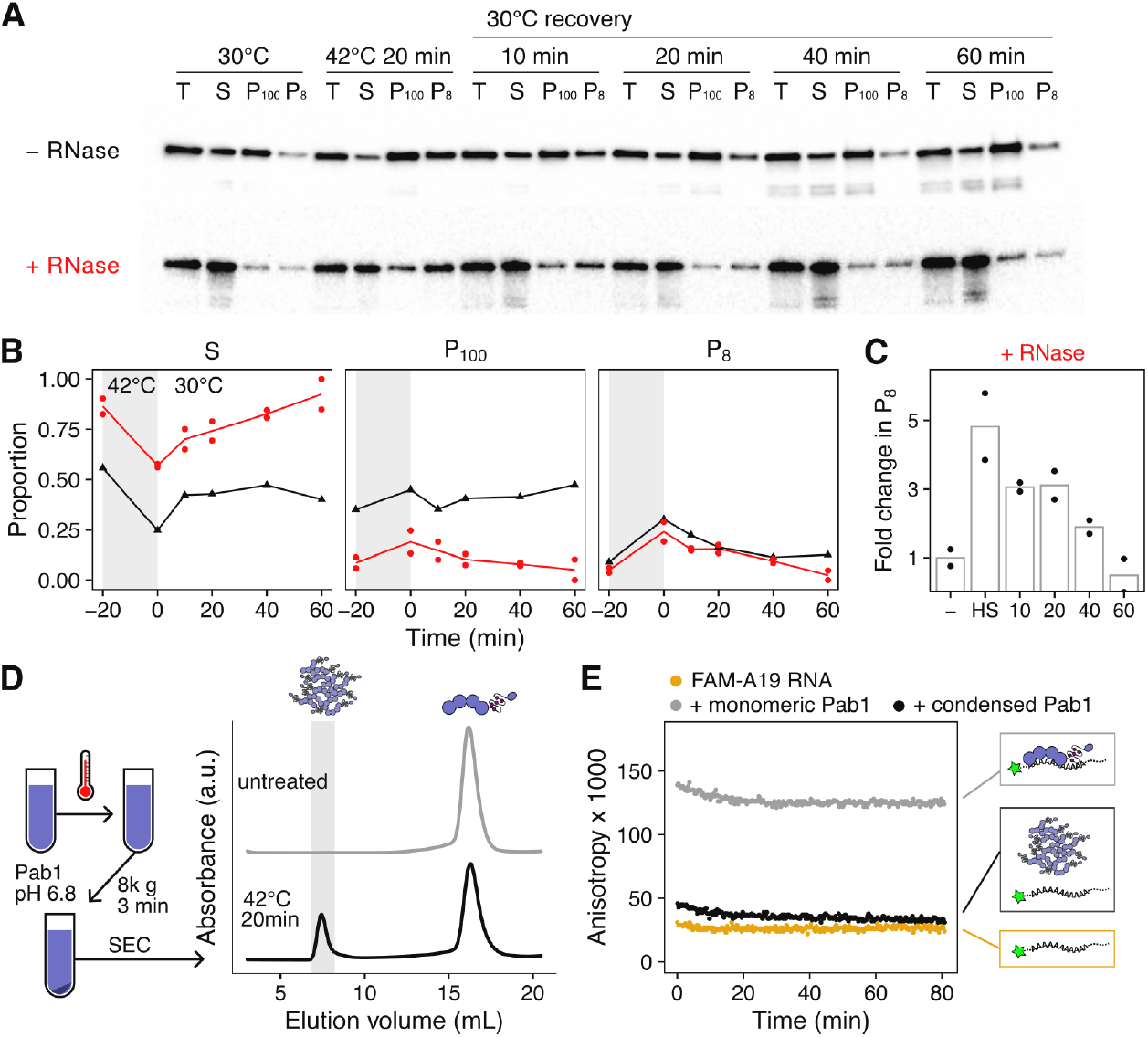
Heat shock causes Pab1 condensation, which is not spontaneously reversible. (A) Western blot against Pab1 isolated from yeast cells before and after 20 minutes of heat shock at 42°C, and during post-stress recovery at 30°C. Cell lysates were incubated with or without RNase I_f_ and centrifuged at 8,000 g and 100,000 g to separate the supernatant (S) from the pellet (P_8_ and P_100_). (B) Quantification of (A). Red and black colors correspond to the RNase- or mock-treated sample, respectively. (C) Relative change in the fraction of large sedimentable Pab1 (P_8_) after heat shock and during recovery compared to pre-shock level. (D) Schematic description of *in vitro* Pab1 condensate purification process and the representative SEC traces for untreated and heat shocked Pab1. Only the heat shocked sample contains Pab1 condensates, which elute in the void volume shaded in gray. (E) Fluorescence anisotropy of 5’ labeled 19-mer poly(A) RNA (A19) in the absence or presence of Pab1. Pab1 condensates have substantially reduced RNA binding capacity than the equimolar amount of monomeric Pab1. No spontaneous reversal of Pab1 condensates was observed at physiological recovery condition (pH 7.3 buffer at 30°C).

To reconstitute Pab1 condensates *in vitro*, we treated purified Pab1 at 42°C for 20 minutes in a physiological buffer at pH 6.8, which is about the measured pH of budding yeast cytoplasm during the same heat shock (Triandafillou et al., 2020) (Figure 1D). We examined the size distribution of Pab1 in the soluble fraction using size exclusion chromatography (SEC) and saw a clear division of Pab1 into two peaks: one peak corresponding to Pab1 monomers and another peak in the void volume corresponding to Pab1 condensates larger than 5,000 kDa (Figure 1D). About 30% of total recombinant Pab1 shifted to the void peak after heat shock, consistent with about 25% of total cellular Pab1 sedimented at 8,000 g spin (Figure 1B). Because a previous study indicated that misfolded proteins can nucleate stress granule formation *in vivo* (Kroschwald et al., 2015), we tested whether heat shocking Pab1 in the presence of firefly luciferase increases the condensation yield. Indeed, we found that heat shocking Pab1 in the presence of 100-fold lower amount of thermolabile firefly luciferase, but not thermostable BSA, increased the condensation yield to about 50% (Figure S1A-C).

Condensation requires Pab1’s folded RNA recognition motifs (RRMs) and excess RNA inhibits Pab1 condensation *in vitro* (Riback et al., 2017), suggesting a competition between Pab1’s condensation and RNA-binding activity. Consistent with this, heat shock reduces the association of Pab1 with RNA *in vivo* (Bresson et al., 2020). We measured the RNA-binding capacity of Pab1 condensates isolated from SEC using fluorescence anisotropy. For 1:1 binding of Pab1 to RNA, we made 19-mer poly(A) RNA (A19) and labeled the 5’ end of the RNA with fluorescein. Indeed, Pab1 condensates showed significantly reduced RNA binding activity compared to monomers (Figure 1E).

As expected, unlike Pab1 condensates formed *in vivo*, Pab1 condensates formed *in vitro* remained stable and RNA-binding incompetent even after dilution into pH 7.3 buffer at 30°C (Figure 1E), consistent with a requirement for cellular disaggregation machinery as repeatedly indirectly demonstrated. Thus, we next investigated whether the reversal of Pab1 condensates to RNA-binding monomers depends on direct engagement the molecular disaggregation system.

### Hsp104, Hsp70, and type II Hsp40 are necessary and sufficient for complete dispersal of Pab1 condensates *in vitro*

To monitor the dispersal Pab1 condensates into functional monomers, we developed a fluorescence anisotropy-based assay in which the increase in fluorescence anisotropy of labeled A19 RNA indicates RNA binding by Pab1 (Figure 2A). We mixed Pab1 condensates and labeled A19 with molecular chaperones Hsp104, Ssa2 (Hsp70), Ydj1 and Sis1 (type I and II Hsp40, respectively), and Sse1 (Hsp110). In the absence of ATP, no change in fluorescence anisotropy was observed. In contrast, in the presence of 5 mM ATP, fluorescence anisotropy quickly increased and reached a plateau after about 5 minutes, marking the completion of Pab1 dispersal (Figure 2A).

**Figure 2.**
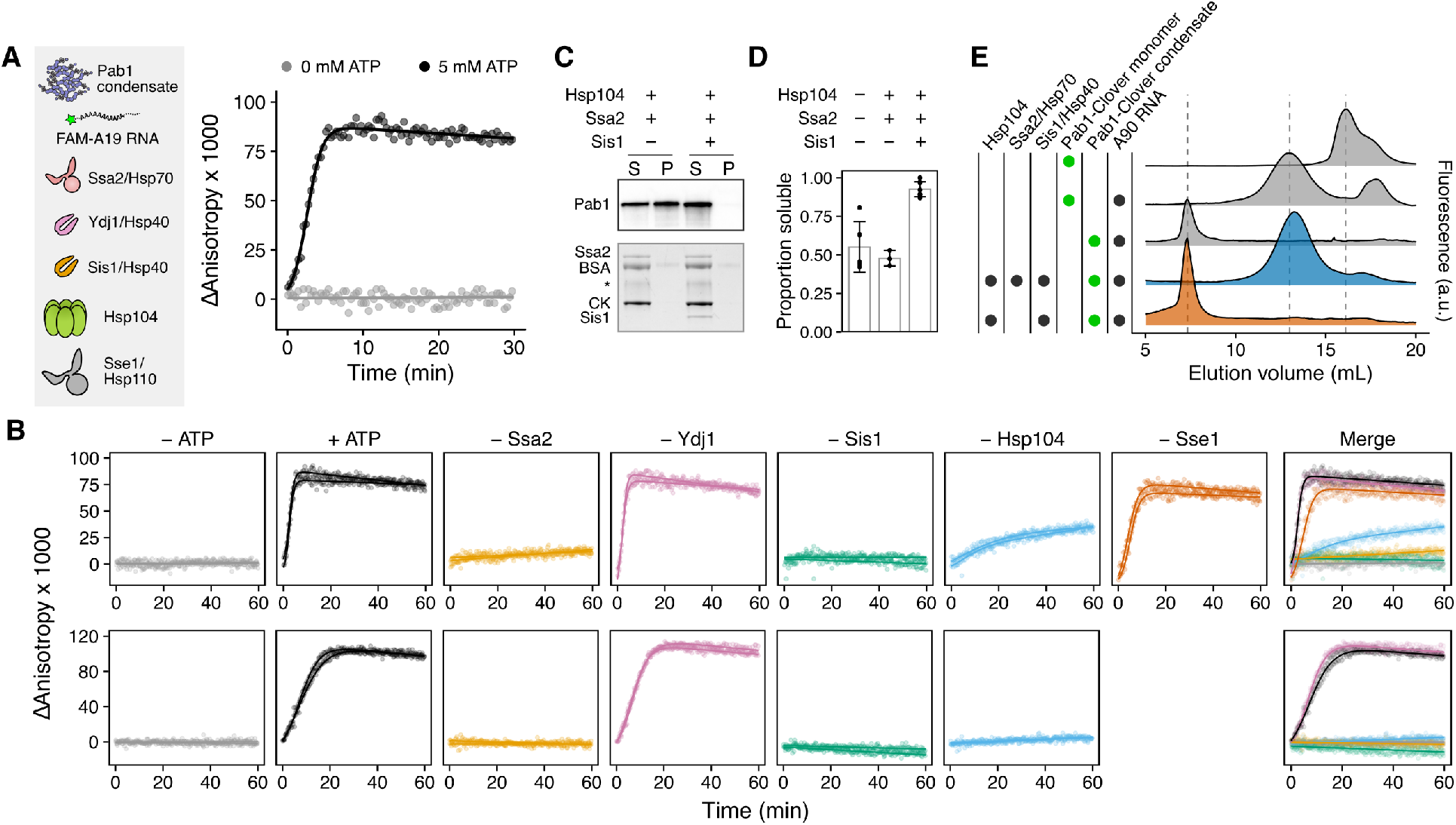
Hsp104, Hsp70, and Sis1 are necessary and sufficient for complete dispersal of Pab1 condensates *in vitro*. (A) Schematic representation of Pab1 condensate, labeled A19 RNA, and molecular chaperones used in the fluorescence anisotropy assay. Solid line is the fit of experimental data points to the logistic equation. (B) Time-resolved fluorescence anisotropy of A19 in the presence of Pab1 condensates and a specific set of molecular chaperones. All chaperones shown in (A) except the component specified at the top of each column were included in the experiments shown in the top row. The same experiment repeated in the absence of Sse1 is shown in the bottom row. Fitted data points from two independent experiments are shown. Merged data points and a solid line fitted to the merged data are shown. (C) Western blot of Pab1 after sedimentation. Total protein image of the corresponding lanes is shown as a loading control at the bottom. CK stands for creatine kinase. Asterisk indicates unknown contaminant. (D) Quantification of Pab1 sedimentation results. (E) Fluorescence-detection SEC (FSEC) profiles of Pab1-Clover. The dashed lines mark the peaks corresponding to Pab1-Clover condensates (7.3 mL), RNA-bound Pab1-Clover monomers (13 mL), and free monomers (16 mL).

We next tested which set of molecular chaperones are necessary and sufficient for complete Pab1 dispersal *in vitro* by removing one component of the chaperone mix at a time and monitoring the effect on Pab1 dispersal (Figure 2B). We used excess molecular chaperones except Sse1, which becomes inhibitory when present in excess (Kaimal et al., 2017), to help ensure even weak disaggregation activity would be detected. Condensate dispersal in the absence of Sse1 absolutely required ATP, Ssa2, Sis1, and Hsp104 (Figure 2B; bottom). When Sse1 was present, removal of Hsp104 led to a much slower and incomplete dispersal of Pab1 (Figure 2B; top). This is consistent with the weak disaggregation activity of Hsp110/70/40 observed against amorphous aggregates and amyloid fibrils (Shorter, 2011). Sse1 and Ydj1 were dispensable in the presence of Hsp104, Ssa2, and Sis1 for both condensates formed in the absence or presence of luciferase (Figure S1D).

The same chaperone requirement pattern was observed when we repeated the assay with Ssa1 and Ssa4, which are respectively the constitutively expressed and heat-inducible paralogs of Ssa2 (Figure S3 and Figure S2A-C). The overall dispersal rate was slower with Ssa4 than with Ssa2, which is consistent with the weaker activity of stress-inducible human Hsp70 observed against amyloid fibrils (Gao et al., 2015; Scior et al., 2018).

We verified our fluorescence anisotropy results using two additional independent methods. First, we examined solubilization of Pab1 using sedimentation. About half of Pab1 condensates isolated from SEC remained in the supernatant after 100,000 g spin in the absence of chaperones, suggesting some condensates are too small to be pelleted (Figure 2C and 2D). Incubating Pab1 condensates with the minimal disaggregation system (Hsp104, Ssa2, Sis1) completely solubilized Pab1. In contrast, Pab1 solubility remained unchanged from background levels when the condensates were incubated with an incomplete disaggregation system.

Next, we prepared Pab1-Clover condensates and examined their size distribution by fluorescence-detection SEC (FSEC). Pab1-Clover condensates remained stable when incubated for an hour at 30°C in the absence of any molecular chaperones or in the presence of an incomplete disaggregation system (Figure 2E). After incubation with the minimal complete disaggregation system, however, the condensate peak disappeared and a new peak corresponding to RNA-bound Pab1-Clover appeared. A similar experiment performed with unlabeled Pab1 using SEC and western blot confirmed that small Pab1 condensates remain stable and are reversed back to monomers only upon incubation with the complete disaggregation system (Figure S3B-C).

In summary, the results from three independent methods consistently indicate that Hsp104, Hsp70, and type II Hsp40 Sis1 are necessary and sufficient for complete dispersal of Pab1 condensates *in vitro*. These results are consistent with the *in vivo* observations that deletion or inhibition of Hsp104, Hsp70, or Hsp40 delays dispersal of Pab1 condensates (Cherkasov et al., 2013) and of stress granules marked by Pab1 (Cherkasov et al., 2013; Walters et al., 2015; Kroschwald et al., 2015, 2018).

### Pab1 condensates and misfolded protein aggregates exhibit different chaperone dependence for dispersal

The mechanism of substrate dispersal by the disaggregation system has been extensively studied using non-native model substrates. The most commonly used model substrate is firefly luciferase, which readily misfolds *in vitro* at elevated temperatures and whose light-producing enzymatic activity can be easily and accurately measured as a readout for the extent of protein refolding. Therefore, we used luciferase as a benchmark heat-misfolded substrate and studied how Pab1 dispersal differs from luciferase disaggregation. We first focused on two chaperone-related features of luciferase disaggregation which we could recapitulate: 1) dependence on co-aggregation with excess Hsp26 for more efficient disaggregation (Cashikar et al., 2005) and 2) synergistic disaggregation in the presence of both type I and II Hsp40s, Ydj1 and Sis1 (Figure 3B) (Nillegoda et al., 2015, 2017).

**Figure 3.**
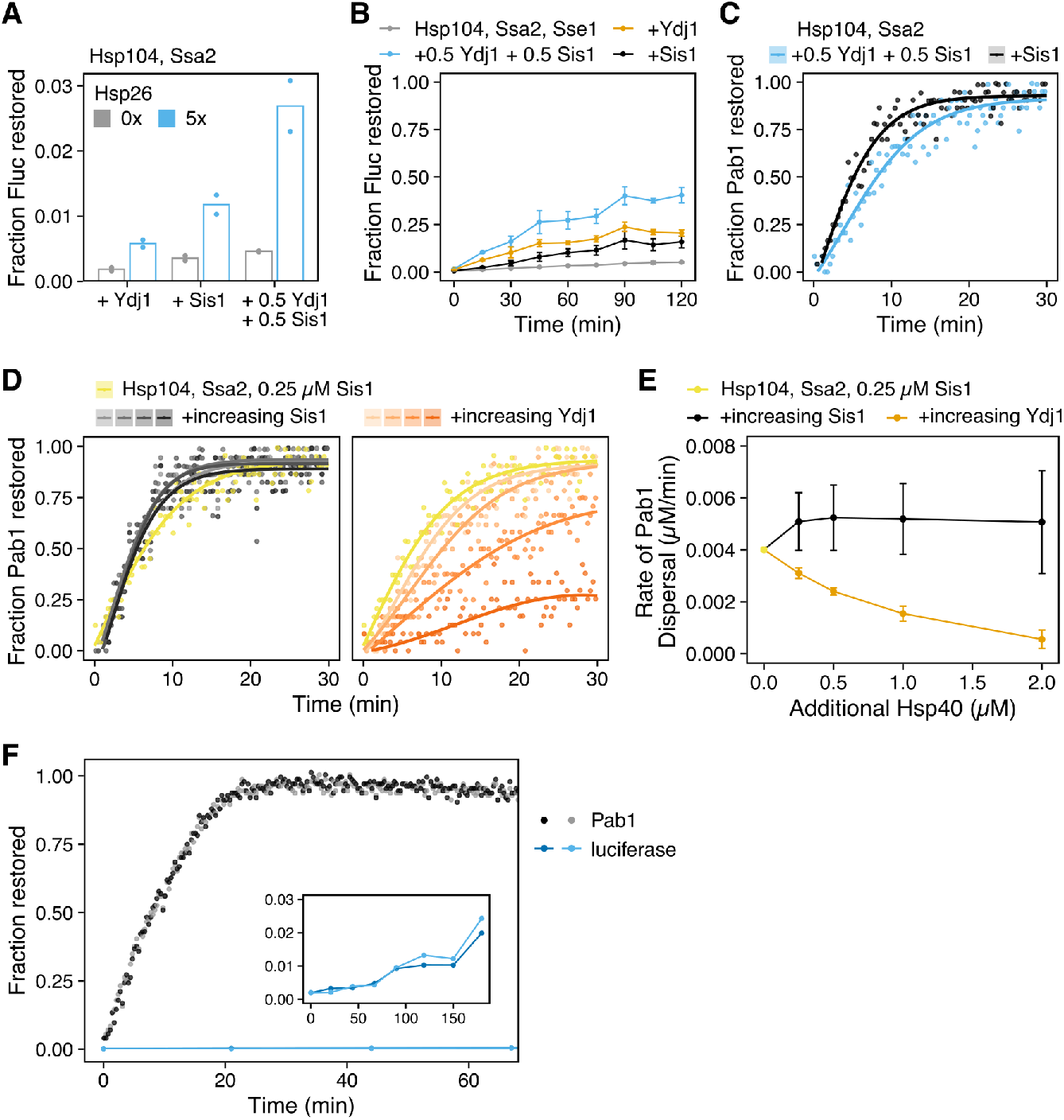
Pab1 condensates and misfolded protein aggregates exhibit different chaperone dependence for dispersal. (A) Fraction of functional luciferase after a two-hour incubation of luciferase aggregates (0.2 µM) with Hsp104 (0.1 µM; 0.5x), Ssa2 (1 µM; 5x), and Hsp40 (0.5 µM total; 2.5x). Luciferase was aggregated either in the absence or presence of 5-fold excess Hsp26. (B) Reactivation of 20 nM aggregated luciferase in the presence of excess Hsp104 (0.75 µM; 37.5x), Ssa2 (0.75 µM; 37.5x) and Sis1 (0.25 µM; 12.5x). Either Sis1 (black), Ydj1 (orange), or both (blue) were used as co-chaperones. Mean and standard deviation were calculated from two independent experiments with duplicates in each experiment. (C) Pab1 dispersal using either Sis1 (black) or a combination of Sis1 and Ydj1 (blue) as co-chaperones. (D) Titration of either Sis1 (left; gradients of black) or Ydj1 (right; gradients of orange) to reactions containing 0.2 µM Pab1 condensates, 0.05 µM Hsp104, 0.5 µM Ssa2, and 0.25 µM Sis1. The amount of additional Hsp40 added was 0.25, 0.5, 1, and 2 µM. (E) The average maximal rate of dispersal and standard deviations quantified from three independent titration experiments, one of which is shown in (D). (F) Restoration of 0.2 µM of Pab1 or luciferase by 0.1 µM Hsp104 (0.5x), 1 µM Ssa2 (5x), and 0.5 µM Sis1 (2.5x). The inset shows zoomed-in refolding kinetics of luciferase over three hours.

Heat-induced aggregation of luciferase in the presence of five-fold excess Hsp26 facilitated subsequent disaggregation and reactivation of luciferase (Figure 3A) as previously reported (Cashikar et al., 2005). To investigate the effect of Hsp26 on Pab1, we subjected Pab1 to a more severe heat shock condition (46°C for 20 minutes at pH 6.4) in the absence or presence of increasing concentrations of Hsp26. Hsp26 suppressed Pab1 condensation and sedimentation in a concentration-dependent manner (Figure S4A-B). Dynamic light scattering (DLS) also revealed that Hsp26 suppresses nucleation of Pab1 (Figure S4C-D). Thus, unlike luciferase which readily co-aggregates with Hsp26 under physiological heat shock conditions, Pab1 condensation is suppressed by Hsp26. Most importantly, unlike luciferase aggregates, Pab1 condensates formed in the absence of Hsp26 are rapidly and completely dispersed by the disaggregation system (Figure 2).

We next investigated whether Pab1 dispersal is accelerated in the presence of both Ydj1 and Sis1, as with luciferase. Ydj1 is a type I Hsp40 which has an highly conserved N-terminal J domain followed by G/F-rich region, zinc-finger domain, C-terminal domains, and a dimerization domain (Kampinga and Craig, 2010). Type II Sis1 largely resembles the architecture of Ydj1 but lacks the zinc-finger domain. To quantify the maximal rate of dispersal in the presence of either or both types of Hsp40s, we converted the fluorescence anisotropy to Pab1 concentration using a calibration curve (Figure S4E and Eq. 3) and extracted the rate (Eqs. 4 and 5). The rate of dispersal did not improve when both Sis1 and Ydj1 were added to Pab1 condensates compared to when only Sis1 was added (Figure 3B-C). Instead, Ydj1 slightly inhibited Pab1 dispersal in a concentration-dependent manner (Figure 3D and 3E). These results indicate that, unlike luciferase aggregates for which Sis1 and Ydj1 show synergistic activity, Sis1 and Yd1j show antagonistic activity for Pab1 condensates.

### The disaggregation system restores Pab1 condensates far more efficiently than misfolded protein aggregates

The poor activity of the disaggregation system against aggregates of model substrates has been observed since the first biochemical reconstitution of the system (Glover and Lindquist, 1998; Goloubinoff et al., 1999). Even with co-aggregation with 5-fold excess Hsp26, less than half of luciferase activity is regained after a two-hour incubation with 37.5-fold excess Hsp104 and Ssa2 (Figure 3B). Indeed, the standard *in vitro* disaggregation protocol requires the use of 10-to 100-fold excess molecular chaperones over substrates to obtain moderate to good yield (Figure 6C). We found that when sub-stoichiometric concentration of Hsp104 (0.5 ×) and closer to stoichiometric concentrations of Ssa2 (5×) and Sis1 (2.5 ×) are used, Pab1 dispersal still completes within 20 minutes while less than 1% of luciferase is reactivated after an hour (Figure 3F). The heat shock condition used to prepare Pab1 condensates and luciferase aggregates was identical except for the initial concentration (2 *µ*M luciferase in the presence of five-fold excess Hsp26 versus 25 *µ*M Pab1).

What causes this large difference in restoration efficiency between Pab1 and luciferase? To gain insight into the potential sources of this discrepancy, we turned to computational kinetic modeling.

### Higher disaggregation rate and partition coefficient lead to more efficient substrate restoration *in silico*

Pab1 condensation requires the folded RRMs, and condensation involves only minor unfolding of secondary structures (Riback et al., 2017). We hypothesized that separation of specific RRM interactions and subsequent folding of the re-solvated Pab1 into native structure may be more efficient compared to the restoration of luciferase aggregates. To test this hypothesis within our model, we synthesized existing simulation studies (Powers et al., 2012; De Los Rios and Barducci, 2014; Nguyen et al., 2017; Xu, 2018; Goloubinoff et al., 2018; Assenza et al., 2019; Wentink et al., 2020) to build what we call a cooperative model of the disaggregation system (Figure 4A and Figure S5A). The cooperative model captures the current model of Hsp104 regulation by Hsp70, in which binding of more than one Hsp70 is required to activate Hsp104 (Seyffer et al., 2012; Carroni et al., 2014). Many of the rate parameters involved, especially in the refolding step, have been measured using bacterial chaperones and model substrates or peptides (Table 1). We assumed that these parameters are generally consistent in the eukaryotic system, and that the same model architecture can be used for both luciferase and Pab1. For details of this ordinary differential equation model, see Methods.

**Figure 4.**
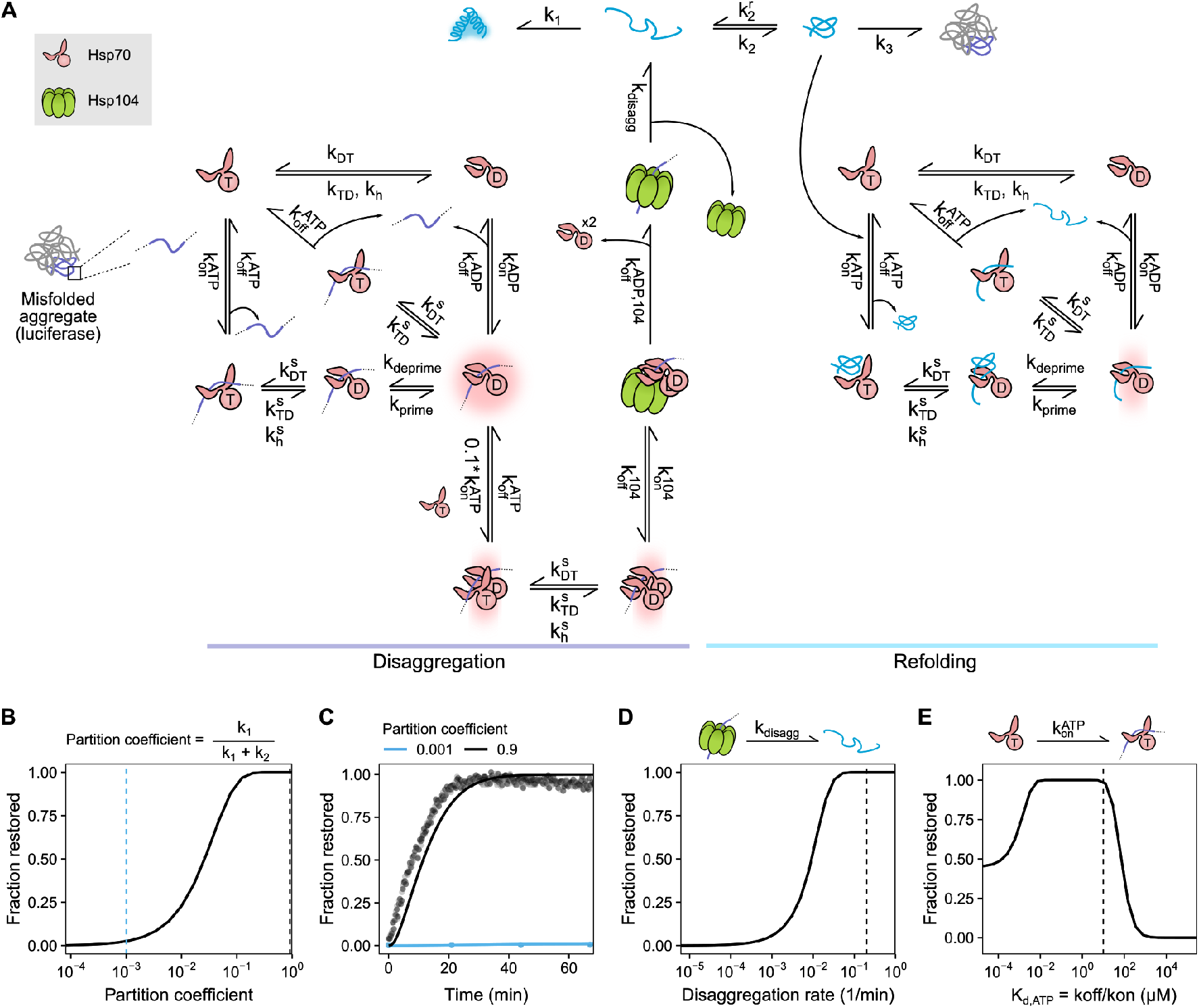
Higher disaggregation rate and folding partition coefficient lead to more efficient substrate restoration *in silico*. (A) Cooperative model of the disaggregation system. For more details, see Methods and Figure S5. (B) Summary of model output in terms of fraction substrate restored as a function of partition coefficient. Dashed lines indicate the partition coefficient used to simulate Pab1 (black) and luciferase (light blue) results in (C). For all simulation experiments in this figure, fraction restored at 2 hours is shown. The starting simulation condition was 0.2 µM substrate, 0.1 µM Hsp104, and 1 µM Hsp70. (C) Simulated substrate dispersal kinetics with either high (black) or low (light blue) partition coefficients. Simulation results (solid line) are overlaid on top of Pab1 and luciferase experimental data from Figure 3F. (D) Simulated fraction substrate restored as a function of disaggregation rate. (E) Simulated fraction substrate restored as a function of Hsp70(ATP) substrate affinity. Dashed lines in (D) and (E) indicate the default value used in the simulation experiments.

**Table 1.** Model parameters.

| Name | Value | Unit | Description | Source |
| --- | --- | --- | --- | --- |
| $k_h$ | 0.036 | $\text{min}^{-1}$ | ATP hydrolysis rate of free Hsp70 | McCarty et al. (1995)* |
| $k_h^s$ | 108 | $\text{min}^{-1}$ | ATP hydrolysis rate of substrate-bound Hsp70 | Laufen et al. (1999)* |
| $k_{\text{on}}^{\text{ATP}}$ | 12 | $\text{min}^{-1} \mu\text{M}^{-1}$ | Substrate onrate to Hsp70<sub>ATP</sub> | Schmid et al. (1994); Gisler et al. (1998)* |
| $k_{\text{off}}^{\text{ATP}}$ | 120 | $\text{min}^{-1}$ | Substrate offrate to Hsp70<sub>ATP</sub> | Schmid et al. (1994); Gisler et al. (1998)* |
| $k_{\text{on}}^{\text{ADP}}$ | 0.06 | $\text{min}^{-1} \mu\text{M}^{-1}$ | Substrate onrate to Hsp70<sub>ADP</sub> | Mayer et al. (2000)* |
| $k_{\text{off}}^{\text{ADP}}$ | 0.0282 | $\text{min}^{-1}$ | Substrate offrate to Hsp70<sub>ADP</sub> | Mayer et al. (2000)* |
| $k_r^{\text{ATP}}$ | 0.008 | $\text{min}^{-1}$ | ATP offrate from Hsp70<sub>ATP</sub> | Russell et al. (1998)* |
| $k^{\text{ATP}}$ | 7.8 | $\text{min}^{-1} \mu\text{M}^{-1}$ | ATP onrate to Hsp70<sub>apo</sub> | Russell et al. (1998)* |
| $k_r^{\text{ADP}}$ | 1.32 | $\text{min}^{-1}$ | ADP offrate from Hsp70<sub>ADP</sub> | Theyssen et al. (1996); Russell et al. (1998)* |
| $k^{\text{ADP}}$ | 16 | $\text{min}^{-1} \mu\text{M}^{-1}$ | ADP onrate to Hsp70<sub>apo</sub> | Russell et al. (1998)* |
| $k_{\text{TD}}, k_{\text{TD}}^s$ | 0.0014 | $\text{min}^{-1}$ | Nucleotide exchange rate from ATP to ADP in free or substrate-bound Hsp70 | Calculated as described in De Los Rios and Barducci (2014) |
| $k_{\text{DT}}, k_{\text{DT}}^s$ | 1.0951 | $\text{min}^{-1}$ | Nucleotide exchange rate from ADP to ATP in free or substrate-bound Hsp70 | Calculated as described in De Los Rios and Barducci (2014) |
| $k_{\text{on}}^{104}$ | 30 | $\text{min}^{-1} \mu\text{M}^{-1}$ | Hsp104 onrate to Hsp70<sub>ADP</sub>:substrate complex | Estimated from the $K_d$ reported in Rosenzweig et al. (2013) |
| $k_{\text{off}}^{104}$ | 60 | $\text{min}^{-1}$ | Hsp104 offrate from Hsp70<sub>ADP</sub>:substrate:Hsp104 complex | Estimated from the $K_d$ reported in Rosenzweig et al. (2013) |
| $k_{\text{off}}^{\text{ADP},104}$ | 100 | $\text{min}^{-1}$ | Hsp70<sub>ADP</sub> offrate after substrate handover to Hsp104 | Free parameter |
| $k_{\text{disagg}}$ | 0.2 | $\text{min}^{-1}$ | Disaggregation rate | Estimated from the measurement in this study |
| $k_{\text{prime}}$ | 10 | $\text{min}^{-1}$ | Substrate remodeling rate | Free parameter |
| $k_{\text{deprime}}$ | 1 | $\text{min}^{-1}$ | Reverse substrate remodeling rate | Free parameter |
| $k_1$ | 10, 0.001 | $\text{min}^{-1}$ | Rate of transition from the unfolded Pab1/luciferase to the folded state | Free parameter |
| $k_2$ | 1 | $\text{min}^{-1}$ | Rate of transition from the unfolded state to the misfolded state | Free parameter |
| $k_2^r$ | 0.1 | $\text{min}^{-1}$ | Rate of transition from the misfolded state to the unfolded state | Free parameter |
| $k_3$ | 1 | $\text{min}^{-1}$ | Rate of transition from the misfolded state to the aggregated state | Free parameter |

\*listed in De Los Rios and Barducci \(2014\) 843

We examined how varying each of the following parameters affected the substrate restoration yield: 1) rate of disaggregation by Hsp104, 2) efficiency of the released substrate from regaining its native structure, which we define as the partition coefficient, and 3) substrate affinity for Hsp70. Modulation of each parameter over 1-2 orders of magnitude substantially affected the restoration yield, measured from 0 to 1 (Figure 4B, D, E). A large difference in partition coefficient alone reproduced the Pab1 and luciferase dispersal data (Figure 4C). The simulation also revealed Hsp70 affinity as a potential factor which can contribute to the observed difference in dispersal efficiency.

Because Hsp104 is associated with both the disaggregation rate and the partition coefficient of a substrate, e.g., through complete threading vs. partial threading of a substrate, we decided to investigate experimentally and compare how Hsp104 engages with Pab1 condensates and luciferase aggregates.

### Pab1 is partially threaded by Hsp104

Substrate threading through the central channel of Hsp104 is a common mechanism for protein disaggregation (Tessarz et al., 2008). Complete threading requires complete substrate unfolding. To probe the folding state of Pab1 and luciferase during their release from Hsp104, we used a mutated version of a bacterial chaperonin, GroEL, which traps unfolded protein (Weber-Ban et al., 1999) (Figure 5A). We verified that this GroEL variant traps unfolded luciferase released during disaggregation and prevents folding (Figure 5B).

**Figure 5.**
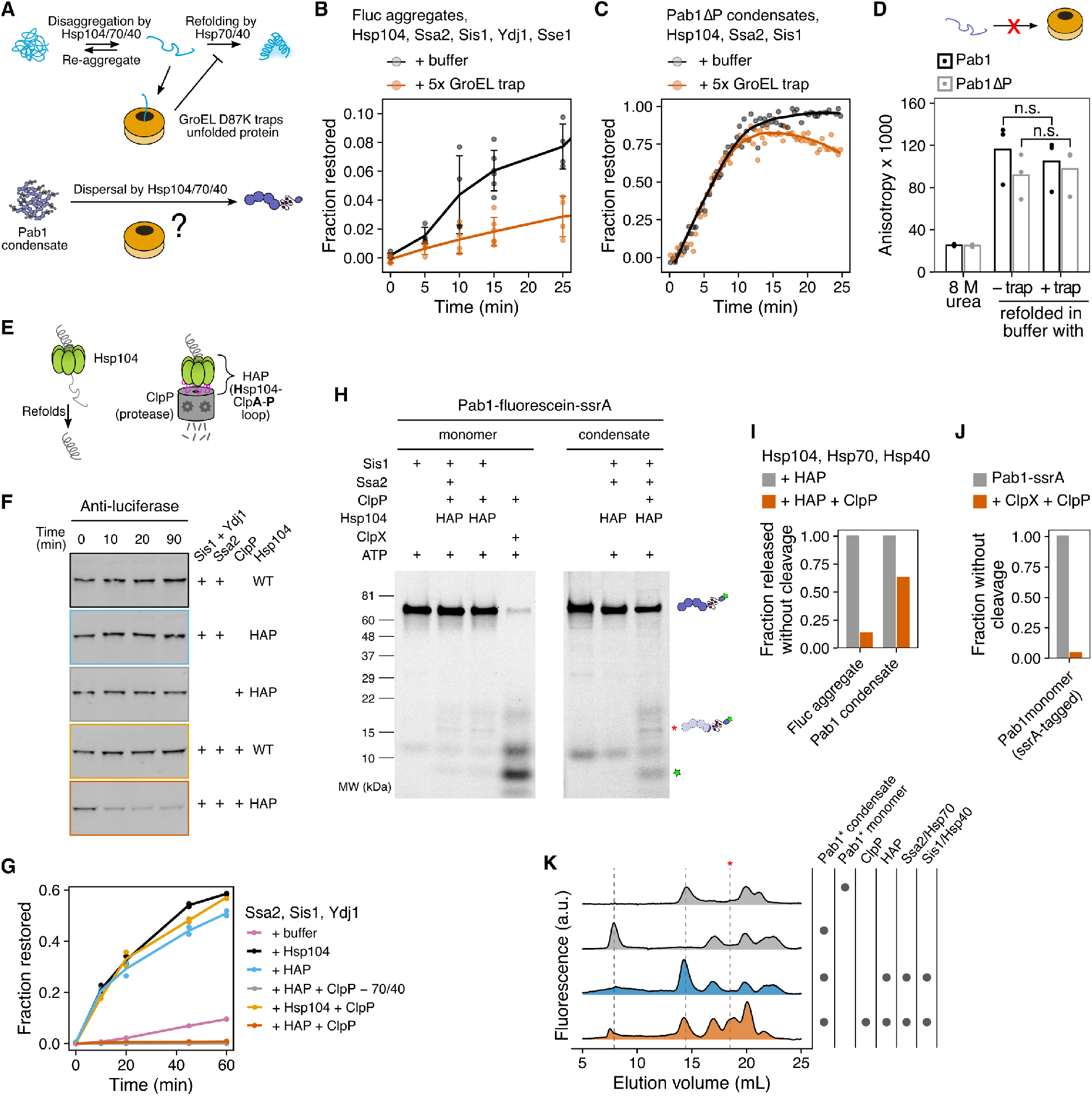
Pab1 is partially threaded by Hsp104. (A) Schematic description of GroEL trap system. (B) Luciferase disaggregation in the absence or presence of 5-fold excess GroEL trap. Mean and standard deviation were calculated from three independent experiments. (C) Pab1ΔP dispersal in the absence or presence of 5-fold excess GroEL trap. Solid lines represent smoothed data from a representative experiment. The decrease in the signal in the presence of GroEL trap after 10 min is due to RNA degradation. (D) Refolding of urea-denatured Pab1 (black) and Pab1ΔP (gray) in buffer containing no or 10-fold excess GroEL trap. (E) Schematic description of HAP/ClpP system. (F) Chemically aggregated luciferase were incubated with the indicated components and the extent of luciferase degradation was visualized by western blot. (G) Refolding kinetics of chemically aggregated luciferase in the presence of indicated components. (H) SDS-PAGE gels of Pab1-fluorescein-ssrA monomers or condensates after an hour incubation with the indicated components. Full-length Pab1 and C-terminal fragments are indicated by the fluorescein band signal. Asterisk indicates HAP/ClpP-specific band. (I) Fraction of full-length protein released from luciferase aggregates or Pab1 condensates, normalized to HAP control. The restoration yield of 60% was used for both luciferase and Pab1 based on the disaggregation and FSEC results, respectively. (J) Fraction of full-length Pab1-ssrA monomers after incubation with ClpX and ClpP. (K) FSEC traces of Pab1-fluorescein-ssrA condensates after an hour incubation with the indicated components. Dashed lines indicate peaks corresponding to Pab1 condensates (8 mL), full-length monomers (14.5 mL), and HAP/ClpP-specific C-terminal fragment (18.5 mL). Unmarked peaks in the control reactions are unknown contaminants.

We then tested whether Pab1 becomes unfolded during dispersal by dispersing Pab1ΔP in the presence of excess GroEL trap. We used Pab1ΔP (55 kDa) because most GroEL substrates are known to have molecular weight less than 60 kDa (Houry et al., 1999) and full-length Pab1 (64 kDa) exceeds that limit. Pab1ΔP lacks the disordered P domain but retains the ability to phase separate (Riback et al., 2017). Pab1ΔP condensates were readily dispersed by the disaggregation system, indicating that the P domain is dispensable for the condensate’s engagement with chaperones (Figure 5C). The rate of dispersal was almost identical for Pab1ΔP dispersed in the absence or presence of GroEL trap. However, when we chemically denatured both full-length Pab1 and Pab1ΔP in 8 M urea and refolded the proteins in buffer containing no or 10-fold excess GroEL trap, both constructs refolded to the same level regardless of the presence of GroEL trap (Figure 5D). This indicated to us that GroEL trap is unable to engage with fully unfolded Pab1 and thus not applicable to investigating the folding state of Pab1.

To circumvent the limitation of the GroEL trap, we turned to the HAP/ClpP system (Tessarz et al., 2008). HAP (Hsp104-ClpA-P loop) is an engineered Hsp104 that interacts with the bacterial peptidase ClpP to form a proteolytic system (Figure 5E). HAP behaved like wild type Hsp104 and quickly degraded luciferase in the presence of ClpP (Figure 5F-G), consistent with a previous report that luciferase is fully threaded and degraded by HAP/ClpP (Haslberger et al., 2008). HAP also behaved like wild type Hsp104 during Pab1 dispersal (Figure S6A-B).

If complete threading of Pab1 were required for condensate dispersal, we would expect to see complete degradation of Pab1 by HAP/ClpP. We made condensates using Pab1-fluorescein-ssrA and examined the degradation pattern after dispersal using SDS-PAGE (Figure 5H and Figure S6C-D) and FSEC (Figure 5K). A mixed group of full-length and degraded Pab1 populations were observed after dispersal. The appearance of full-length Pab1 monomers suggested partial translocation of Pab1 by HAP and release before Pab1 enters the proteolytic chamber of ClpP (Figure 5I and Figure 5K). We confirmed that ClpP can degrade Pab1 using ClpX, which recognizes the ssrA degradation tag and unfolds the substrate for ClpP (Figure 5H and Figure 5J). Specific degradation fragments appeared upon incubation of HAP/ClpP and chaperones with Pab1 condensates, but also to a lesser extent with Pab1 monomers, suggesting a basal level of interaction between Pab1 and HAP/ClpP (Figure 5H). Similar C-terminal fragments containing a part of the P domain and the C-terminal domain of Pab1 appeared for Pab1-Clover and Pab1-fluorescein without the ssrA tag (Figure S6E and Figure S6H). However, much less full-length monomer appeared for Pab1-Clover than Pab1-fluorescein-ssrA (Figure S6F), suggesting that a fluorescent label can affect the processing by HAP/ClpP. We also examined the N-terminal fragments using fluorescein-Pab1 and saw appearance of both large fragments of the RRMs and small peptides (Figure S6G). HAP/ClpP fails to completely disperse condensates of Pab1-Clover (Figure S6F), which limited our ability to quantify the fraction of Pab1 released without cleavage for these constructs.

Together, these results show that, unlike luciferase which requires complete threading and unfolding by HAP for disaggregation, partial threading of Pab1 still leads to condensate dispersal. This is consistent with the partial threading mechanism proposed for proteins with a mixture of misfolded and folded domains (Haslberger et al., 2008; Sweeny and Shorter, 2016) and the lack of major secondary structure changes in Pab1 at condensation temperature (Riback et al., 2017). However, we cannot rule out the possibility that wild type Hsp104 processes unlabeled Pab1 differently from what we observed with HAP and labeled Pab1 constructs.

### Cooperative binding of Hsp70 targets condensates for dispersal

How does the disaggregation system recognize Pab1 condensates? To address this question, we first performed a series of fluorescence anisotropy Pab1 dispersal assays with varying chaperone concentrations and quantified the maximal rate of dispersal (Figure 6A and 6B). Pab1 condensate dispersal was most robust to the Hsp104 concentration, showing half-maximal dispersal rate at 1:10 Hsp104:Pab1 ratio. Excess Sse1 was inhibitory and Sse1 worked most optimally at sub-stoichiometric level, consistent with previous observations (Kaimal et al., 2017; Wentink et al., 2020). Pab1 condensate dispersal was most sensitive to the concentrations of Sis1 and Ssa2. In particular, the rate of dispersal plummeted as the Ssa2 concentration approached the stoichiometric level (Figure 6A).

Indeed, the disaggregation system’s dependence on super-stoichiometric Hsp70 for optimal activity has been a long-standing puzzle. Earlier studies investigating this problem with a reconstituted bacterial disaggregation system found that DnaK (bacterial Hsp70) has to be present in excess for maximal disaggregation yield (Goloubinoff et al., 1999; Ben-Zvi et al., 2004). We surveyed *in vitro* disaggregation studies in the literature and found that this dependence on excess Hsp70 is widespread across studies, precise conditions, and substrates (Figure 6C), and our results are no exception.

We decided to investigate why excess Hsp70 over substrate is needed for what is still a catalytic series of reactions. We titrated Ssa2 over a narrow window around the stoichiometric Hsp70:Pab1 ratio and monitored Pab1 dispersal reaction for eight hours (Figure 6E). We also simulated the cooperative model (Figure 4A) using the same chaperone concentrations used in the *in vitro* experiment (Figure 6E). The cooperative model recapitulated the disaggregation system’s Hsp70-sensitive Pab1 dispersal activity (Figure 6F). This model reflects the results from recent studies which indicate that interaction with more than one Hsp70 is required for activation of Hsp104 (Carroni et al., 2014; Seyffer et al., 2012). Indeed, simulation of non-cooperative model, in which single Hsp70 is sufficient to recruit and activate Hsp104, resulted in high Pab1 dispersal activity even with sub-stoichiometric Hsp70 (Figure 6D and 6F).

The cooperative model was also able to recapitulate the general trend seen in the Hsp104 and Ssa2 titration experiments (Figure 6A and Figure S5B). Although Sis1 and Sse1 were not explicitly included in the model, modulating the ATP hydrolysis rate and ADP exchange rate allowed us to mimic the effect of titrating Sis1 and Sse1, respectively (Figure S5B). Interestingly, although we were able to recapitulate the inhibitory effect of Sse1 with high ADP exchange rate, modulating ADP exchange rate was not enough to recapitulate the facilitative effect of sub-stoichiometric Sse1 (Figure 6G and Figure 6H).

These results converge on a picture in which the presence of multiple, closely spaced Hsp70 molecules on the surface of condensates provide a molecular marker labeling condensates for Hsp104-dependent dispersal, as proposed in the bacterial disaggregation system by Seyffer et al. (2012). Our simulation results indicate that a cooperative Hsp70 effect on Hsp104 binding and activation suffices to explain the disaggregation system’s intrinsic sensitivity to the level of Hsp70.

## Discussion

We demonstrated that the yeast disaggregation system composed of molecular chaperones Hsp104, Hsp70, and Hsp40 can directly reverse heat-induced biomolecular condensates of Pab1 back to functional monomers *in vitro*. This establishes heat-induced biomolecular condensates of Pab1 as endogenous substrates of the molecular disaggregation system. Through comparative studies of Pab1 and the model substrate firefly luciferase, we uncovered a number of distinctions in the way chaperones engage with each substrate. The most notable distinction was the large difference in the restoration efficiency between Pab1 condensates and luciferase aggregates. Our results show that partial threading of Pab1 by Hsp104 during dispersal can contribute to this large difference in the restoration efficiency. Finally, we find that efficient Pab1 dispersal depends on the presence of excess Hsp70, which serves as a condensate detector through cooperative recruitment and activation of Hsp104 near the surface of condensates (Figure 7).

**Figure 6.**
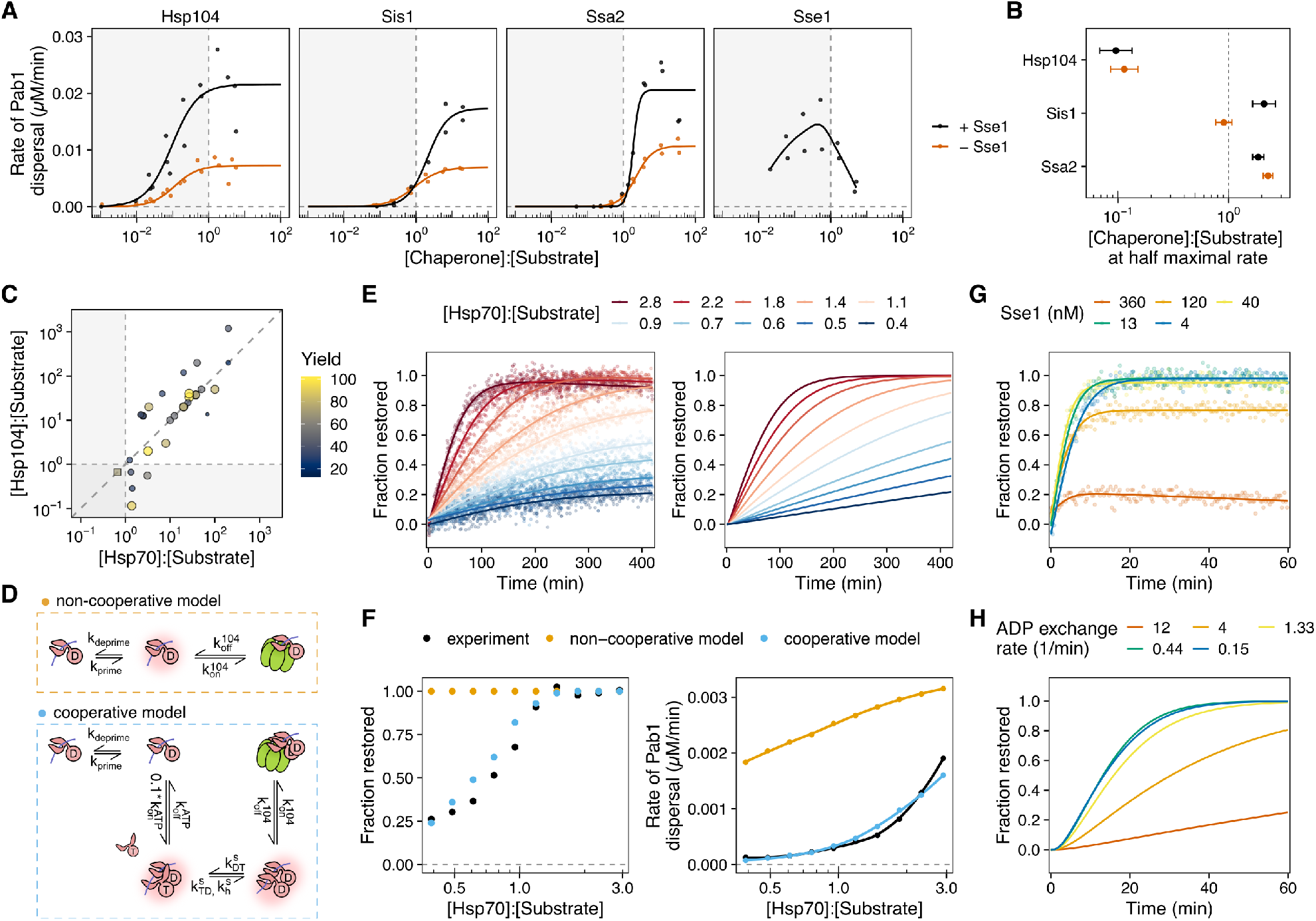
Cooperative binding of Hsp70 labels condensates for disaggregation. (A) Maximal rate of Pab1 dispersal in the presence (black) or absence (brown) of Sse1. Solid lines in Hsp104, Sis1, and Ssa2 panels represent logistic fit to the data. Solid line in Sse1 panel is the smoothing line. The baseline concentrations of the proteins were 0.2 *µ*M Pab1, 0.5 *µ*M Ssa2, 0.5 *µ*M Sis1, 0.2 *µ*M Hsp104, and 0.1 *µ*M Sse1. Shaded area denotes the sub-stoichiometric region. The lower rate with the highest Hsp70 concentration is due to RNA degradation. (B) Relative chaperone concentration at half-maximal dispersal rate, extracted from the fits in (A). Error bars represent standard errors around the estimated parameter. (C) Survey of disaggregation studies in the literature. Maximal yield of disaggregation experiments and the relative amount of Hsp104 and Hsp70 used in the experiment are shown. Color and size of each data point correspond to the yield. Circles represent studies with wild type Hsp104. One square data point in the sub-stoichiometric Hsp70 area represents a study which used a hyperactive D484K variant of Hsp104. (D) Schematic comparison of the cooperative and non-cooperative models. (E) Pab1 dispersal monitored by fluorescence anisotropy (left) and the simulated Pab1 dispersal results from the cooperative model (right). (F) Quantitative comparison of Pab1 dispersal data (black) to the simulated results from the cooperative (blue) and non-cooperative models (orange). Fraction restored at the end of the 8 hour experiment (left) and the maximal rate of dispersal (right) were used for comparison. (G) Representative Sse1 titration Pab1 dispersal data. (H) Simulation of Pab1 dispersal with different ADP nucleotide exchange rates using the cooperative model.

**Figure 7.**
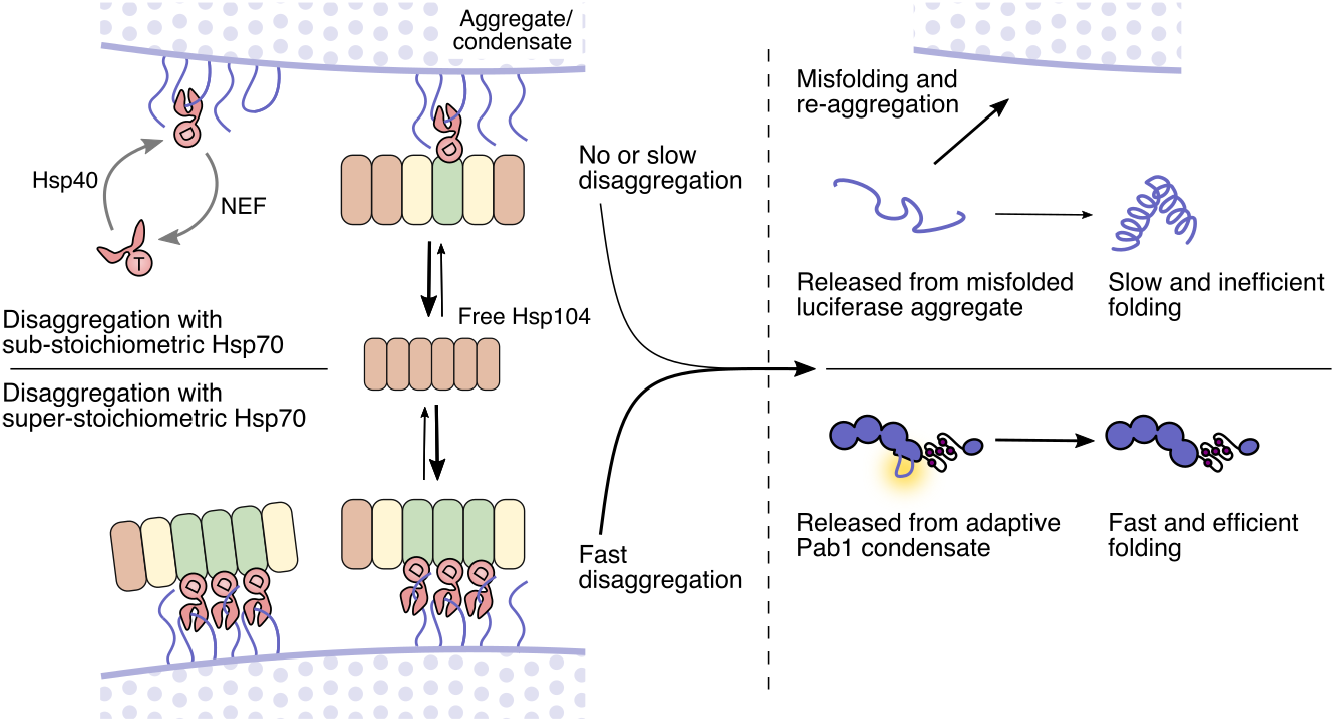
Model of Pab1 dispersal by the Hsp104/Hsp70/Hsp40 disaggregation system. Productive recruitment and activation of Hsp104 depend on the presence of multiple closely spaced Hsp70 molecules on the surface of aggregate/condensate, a condition which is most likely to be achieved with excess Hsp70. Luciferase is fully threaded by Hsp104 and released as unfolded protein; This leads to slow and inefficient folding because unfolded protein is prone to misfolding and re-aggregation. In contrast, Pab1 is released after partial threading by Hsp104 and readily re-folds into native structure, resulting in faster and more efficient folding compared to luciferase. The schematic of Hsp104 with repressed (red) and activated (green) protomers is adapted from Carroni et al. (2014).

### Heat-induced biomolecular condensates are endogenous substrates of the molecular disaggregation system

In yeast, heat-induced biomolecular condensates including stress granules adopt a solid-like characteristic (Kroschwald et al., 2015; Riback et al., 2017). Timely dispersal of endogenous condensates depends on molecular chaperones (Cherkasov et al., 2013; Walters et al., 2015; Kroschwald et al., 2015, 2018), and the timing of dispersal correlates with resumption of active cellular translation and growth (Cherkasov et al., 2013; Kroschwald et al., 2015, 2018). These *in vivo* observations strongly suggest heat-induced biomolecular condensates are the endogenous substrates of the molecular disaggregation system. However, direct biochemical evidence for chaperone-mediated condensate dispersal has been missing.

In this study, we showed using *in vitro* reconstitution that the Hsp104/70/40 disaggregation system directly engages and disperses heat-induced condensates of Pab1. The contrast between Pab1 condensates and luciferase aggregates on multiple important dimensions (requirements for specific chaperones and for small heat-shock proteins, rate and yield of dispersal, the nature of engagement with Hsp104) demonstrates that luciferase, and by extension other similarly behaving “model” misfolded proteins, have severe limitations as models of endogenous heat-induced chaperone substrates. Whether the authentic substrate Pab1 is itself a suitable model for other endogenous substrates remains an important open question.

We also showed that Hsp110, Hsp70, and Hsp40 can disperse Pab1 condensates. This alternative disaggregation system is also capable of disaggregating luciferase aggregates and amyloid fibrils, although with much weaker disaggregation activity than with Hsp104 (Shorter, 2011). The disaggregation activity of the Hsp110/70/40 system is conserved in animals (Shorter, 2011; Wentink et al., 2020), which lack cytosolic Hsp104 (Erives and Fassler, 2015), suggesting a potential evolutionary and biochemical bridge to chaperone-mediated condensate dispersal in animals.

### Different engagement of molecular chaperones with biomolecular condensates and misfolded protein aggregates

Co-aggregation of luciferase with Hsp26 keeps luciferase in a near-native conformation, generates smaller aggregates, and facilitates aggregate interaction with Hsp70 (Cashikar et al., 2005; Ungelenk et al., 2016; Żwirowski et al., 2017). However, Hsp26 is almost undetectable in cells pre-stress (Cashikar et al., 2005). Because biomolecular condensation of endogenous proteins occurs within a few minutes of heat stress (Wallace et al., 2015), Hsp26 is unlikely to be involved in condensate regulation during the initial exposure to stress. We showed that Hsp26 prevents condensation of Pab1 *in vitro*. Human small heat shock protein Hsp27 has also been shown to repress phase separation of FUS (Liu et al., 2020). Post-stress accumulation of Hsp26 in cells may be involved in desensitization of the cells to sustained or repeated stress by modulating the phase boundary of endogenous mature proteins.

The canonical type I (Ydj1; DNAJA2 in humans) and type II Hsp40 (Sis1; DNAJB1) chaperones show synergistic activity toward luciferase aggregates (Nillegoda et al., 2015, 2017). This synergistic activity stems from the preference of type I and II Hsp40 chaperones for small (200-700 kDa) and large (*>*5,000 kDa) aggregates, respectively (Nillegoda et al., 2017), although how aggregates of different sizes can be distinguished at the molecular level remains unclear. Sis1 and Ydj1 also exhibit different amino acid sequence preference (Jiang et al., 2019) and different mode of binding Hsp70, the latter of which has been shown to be responsible for amyloid disaggregation activity unique to type II chaperones (Faust et al., 2020). The inability of Ydj1 to support Pab1 dispersal could be due to the large size of Pab1 condensates, the lack of Ydj1 binding sites among the exposed region of Pab1 in condensates, dependence on the Hsp70 binding mode unique to type II Hsp40, or any combination of these.

The antagonistic effect of Ydj1 on Sis1 suggests a competition between the two co-chaperones for Hsp70. In cells, stress-induced phosphorylation of Hsp70 reprograms Hsp70’s substrate specificity, e.g., by preventing Hsp70 from interacting with Ydj1 but not with Sis1 (Truman et al., 2012). Similar post-translational modifications may also regulate the activity of the disaggregation system toward specific substrates.

Hsp104 functions by threading substrate through its central channel (Gates et al., 2017). However, both our simulation and biochemical experiments suggest Pab1 is partially threaded by Hsp104. A partial threading activity of Hsp104 and its bacterial homolog ClpB has been reported previously, where both proteins selectively thread misfolded moiety of the substrate while leaving the natively folded domains intact (Haslberger et al., 2008; Sweeny and Shorter, 2016). We hypothesize that partial threading of the locally “unfolded” region of Pab1, possibly the same or near the region mediating Pab1 condensation interaction, allows for condensate dispersal without substantial protein unfolding.

### Hsp70 clusters are a potential condensate-specific marker for Hsp104

Efficient dispersal of Pab1 condensates depends on the presence of excess Hsp70. Nearby Hsp70s, which would be rare on monomers but common on condensates, both increase Hsp104 binding and stimulate additional Hsp104 activity. In this sense, consistent with an insightful proposal from Seyffer et al. (2012) working in the homologous bacterial system, Hsp70 clusters provide an active label for engagement and activation of powerful dispersal machinery only in spatial proximity to condensed substrates.

Cooperative action of Hsp70 in substrate unfolding has been proposed to explain the requirement of excess Hsp70 during glucose-6-phosphate dehydrogenase (G6PDH) disaggregation (Ben-Zvi et al., 2004). We find by simulation that cooperative action of Hsp70 in the recruitment and activation of Hsp104 is sufficient to reproduce the *in vitro* Hsp70 titration data. We also found that, in the cooperative model, modulating the ADP exchange rate alone was not enough to reproduce the facilitative effect of Sse1 (Figure 6H). A recent study by Wentink et al. (2020) uncovered an additional function of human Hsp110 in promoting local clustering of Hsp70 on the substrate surface. A similar function in the yeast Hsp110, Sse1/2, may explain the discrepancy between our model and the data.

### Biomolecular condensates in the cellular heat shock response

Engagement of molecular chaperones, especially Hsp40 and Hsp70, with stress-induced biomolecular condensates provides a tangible means to explain how yeast cells integrate multiple physical cues from the environment to sense temperature. In yeast, the transcriptional heat shock response is triggered when Hsp70, which is bound repressively to the transcriptional factor Hsf1, is titrated away by stress-induced substrates (Zheng et al., 2016; Krakowiak et al., 2018; Peffer et al., 2019; Masser et al., 2019; Feder et al., 2021).

*S. cerevisiae* cultured at 30°C begins mounting the transcriptional heat shock response when the temperature is raised above 36°C. The identities of these stress-induced substrates remain elusive. Although misfolding-prone nascent or newly synthesized polypeptides are known to help trigger the response, suppression of protein synthesis is not sufficient to suppress the transcriptional response, implying the existence of mature substrates which, we recently showed, also like depend on stress-associated intracellular acidification for formation (Triandafillou et al., 2020). Notably, condensation of Pab1 and other heat-sensitive proteins is strongly pH-sensitive (Riback et al., 2017; Iserman et al., 2020; Kroschwald et al., 2018). Here, we provide an additional important piece of circumstantial evidence for the model of stress-triggered condensation as an activator of Hsf1: heat-induced condensates of Pab1 are authentic chaperone substrates which depend on Hsp70 for dispersal. Thus, Pab1 can autonomously transduce physiological heat shock temperatures into biomolecular condensation, dependent on pH, and recruit molecular chaperones including Hsp70. In short, Pab1—and by extension presumably others of the dozens of previously identified heat-condensing proteins (Wallace et al., 2015; Cherkasov et al., 2015), including more than a dozen which condense in response to a 37°C heat shock—now appears to have all the characteristics needed to act as an inducer of the transcriptional heat shock response. An important conceptual difference is that while the proteotoxicity model has invoked toxic misfolding, biomolecular condensation is known to be an evolved, adaptive response (Riback et al., 2017; Iserman et al., 2020).

### Molecular chaperones as biomolecular condensate remodelers

In this work, we show that molecular chaperones can regulate biomolecular condensates by acting as dispersal factors. This expands the list of known condensate dispersal factors, which currently includes the dual specificity kinase DYRK3 (Wippich et al., 2013) and nuclear-import receptor karyopherin-*β*2 (Guo et al., 2018). The functional repertoire of molecular chaperones in biomolecular condensate regulation is likely to be much broader than just dispersal. For example, Hsp104, Hsp70, and Hsp40 in yeast are required for condensate formation of SNF1 kinase activator Std1 during fermentation (Simpson-Lavy et al., 2017). Another example is Hsp27, which partitions into liquid-like FUS condensates upon stress-induced phosphorylation and prevents amyloid transition of FUS (Liu et al., 2020). Illumination of the roles of molecular chaperones as facilitators, remodelers, and dispersers of biomolecular condensates—and the mechanisms and biological consequences of this regulation—presents an enormous opportunity for expanding our understanding of these ancient molecules.

## Methods

### Resource Availability

#### Lead Contact

Further information and requests for resources and reagents should be directed to and will be fulfilled by Lead Contact, D. Allan Drummond.

### Materials Availability

All unique and stable reagents generated in this study are available from the Lead Contact.

### Data and Code Availability

#### Experimental data and code for analysis

All data analysis and visualization were performed with R (version 3.5.2) in RStudio (RStudio Team, 2018). The raw and processed data, and the custom scripts for data process, data analysis, and figure generation will be available on Data Dryad.

#### Code for simulation and data analysis

Simulation was performed with Python (version 3.7.7) in Jupyter notebook (Kluyver et al., 2016). The code is be available on GitHub (https://github.com/haneulyoo/sim_disagg_2021).

### Experimental Model and Subject Details

#### Yeast strain and growth conditions

*S. cerevisiae* strain BY4741 (MATa ura3Δ0 leu2Δ0 his3Δ1 met15Δ0) cells were cultured in yeast extract peptone dextrose (YPD) media in shaking baffled flasks at 30°C. The strain background used was S288C.

#### Bacteria strain and growth conditions

Unless specified otherwise under Chemicals, Peptides, and Recombinant proteins of the Key Resources Table, all recombinant proteins used in this work are expressed in and purified from *E. coli* BL21(DE3). Cells were first grown in Luria broth (LB) at 37°C for 12 to 16 hours and then inoculated to 1-2 L Terrific Broth (TB) culture. Specific growth condition used for each recombinant protein is described in Method Details.

## Method Details

### Purification of Pab1 and Pab1 variants

Protein expression and purification protocols were adapted with modification from (Riback et al., 2017). N-terminally 8xHis-tagged Pab1 constructs were transformed into an *E. coli* strain BL21(DE3) and grown overnight at 37°C. The overnight culture was used to inoculate 1L Terrific Broth (TB). Cells were grown until the optical density at 600 nm (OD_600_) reached between 0.4 and 0.6 and then the flask was moved into a 30°C incubator. After 30 minutes, 0.2 mM IPTG was added to induce protein expression. Cells were harvested after 4 hours and lysed by sonication in His binding buffer (20 mM HEPES pH 7.3, 150 mM KCl, 2.5 mM MgCl_2_, 20 mM imidazole, 10 % glycerol, and 1 mM BME) supplemented with 1 Pierce protease inhibitor tablet (Thermo Fisher A32965). Lysate was cleared by spinning at 20,000 g for 15 minutes. Cleared lysate was loaded onto a 5 mL HisTrap FF column (Thermo Fisher 17525501) equilibrated with His binding buffer on an AKTA FPLC system. Protein was eluted with a 20 mL gradient from 0 to 100 % His elution buffer (20 mM HEPES pH 7.3, 150 mM KCl, 2.5 mM MgCl_2_, 400 mM imidazole, 10 % glycerol, and 1 mM BME). Fractions containing Pab1 were buffer exchanged into a Q binding buffer (20 mM HEPES pH 7.3, 50 mM KCl, 2.5 mM MgCl_2_, 10 % glycerol, and 1 mM DTT) and loaded onto a 5 mL HiTrap Heparin HP column (GE Healthcare 17040701) to remove nucleic acids. Nucleic acid-free protein was eluted over a 20 mL gradient from 0 to 100 % Q elution buffer (20 mM HEPES pH 7.3, 1 M KCl, 2.5 mM MgCl_2_, 10 % glycerol, 1 mM DTT). Fractions of interest were combined with an aliquot of a homemade tobacco etch virus (TEV) protease and dialyzed against 1 L His binding buffer overnight to remove the N-terminal tag and to lower the salt concentration. On the next day, the dialyzed solution was loaded again onto a 5 mL HisTrap FF column and the flow-through which contains the cleaved protein was collected. The protein was concentrated and loaded onto a Superose 6 10/300 GL column (GE Healthcare) equilibrated with SEC/Storage buffer (20 mM HEPES pH 7.3, 150 mM KCl, 2.5 mM MgCl_2_, and 1 mM DTT). Monomeric proteins were pooled together, further concentrated if necessary, and stored at −80°C. Protein concentration was measured using Bradford assay (Bio-Rad 5000201).

### Purification of Hsp70 chaperones

We adapted with minor modifications the protocol provided by Zachary March in James Shorter’s group. N-terminally 6xHis-SUMO-tagged Hsp70 proteins were transformed into an *E. coli* strain BL21(DE3) and grown overnight at 37°C. The overnight culture was used to inoculate 1L Terrific Broth (TB). Cells were grown until OD_600_ between 0.4 and 0.6 and then the flask was moved into a 18°C incubator. After 30 minutes, 0.2 mM IPTG was added to induce protein expression overnight. Cells were harvested after 14 - 18 hours and lysed by sonication in Hsp70 His binding buffer (50 mM HEPES pH 7.3, 750 mM KCl, 5 mM MgCl_2_, 20 mM imidazole, 10 % glycerol, 1 mM BME, and 1 mM ATP) supplemented with 1 Pierce protease inhibitor tablet. Cleared lysate was loaded onto a 5 mL HisTrap FF column equilibrated with Hsp70 His binding buffer on an AKTA FPLC system. After loading, the column was washed with more Hsp70 His binding buffer until the UV reading returned to a steady, baseline level. The column was further washed with about 20 mL high ATP buffer (50 mM HEPES pH 7.3, 750 mM KCl, 5 mM MgCl_2_, 20 mM imidazole, 10 % glycerol, 1 mM BME, and 20 mM ATP) and incubated in this buffer for at least 30 minutes. The high ATP buffer was washed out with Hsp70 His binding buffer and the protein was eluted with a 20 mL gradient from 0 to 100 % Hsp70 His elution buffer (50 mM HEPES pH 7.3, 750 mM KCl, 5 mM MgCl_2_, 400 mM imidazole, 10 % glycerol, 1 mM BME, and 1 mM ATP). The fractions of interest were combined and dialyzed against 1 L Hsp70 His binding buffer for at least 2 hours to remove excess imidazole. An aliquot of homemade SUMO protease Ulp1 was added to the dialysis bag. Dialysis was continued overnight at 4°C. Next day, the cleaved protein was recovered by running the dialyzed solution through His column and collecting flow-through. Flow-through fractions containing tag-free Hsp70 proteins were combined, diluted in Hsp70 Q binding buffer (20 mM HEPES pH 7.3, 50 mM KCl, 5 mM MgCl_2_, 0.5 mM EDTA, 2 mM DTT, and 1 mM ATP), and loaded onto an equilibrated 5 mL HiTrap Q HP anion exchange column (GE Healthcare 17115401). Hsp70 was eluted over a 50 mL gradient from 0 to 100 % Hsp70 Q elution buffer (20 mM HEPES pH 7.3, 1 M KCl, 5 mM MgCl_2_, 0.5 mM EDTA, 2 mM DTT, and 1 mM ATP). Fractions containing Hsp70 were determined by SDS-PAGE. We observed a peak with a left shoulder or two closely overlapping peaks around 25 mS/cm. Both peaks contained Hsp70, but only the later peak fractions exhibited activity in both luciferase and Pab1 disaggregation assays. We combined the fractions corresponding to the second peak, concentrated, and buffer exchanged the protein into Hsp70 storage buffer (50 mM HEPES pH 7.3, 150 mM KCl, 5 mM MgCl_2_, 10 % glycerol, 2 mM DTT, and 1 mM ATP). Protein concentration was measured using Bradford assay. Protein aliquots were snap-frozen in liquid nitrogen and stored at −80°C.

### Purification of sortase A enzymes

Wild-type (Guimaraes et al., 2013) (used for N-terminal labeling) and heptamutant sortase A (Hirakawa et al., 2015) (used for C-terminal labeling) were purified using the same protocol. Constructs were transformed into *E. coli* strain BL21(DE3), grown in 1 L of TB until they reached an OD_600_ of 0.6. Protein production was induced with 0.5 mM IPTG. The cells were incubated overnight at 18°C, and harvested in His binding buffer (20 mM HEPES pH 7.5, 150 mM KCl, 2.5 mM MgCl_2_, 20 mM imidazole, 10 % glycerol, and 1 mM BME), supplemented with protease inhibitors and Pierce Universal Nuclease (Thermo Scientific PI88702). Cells were lysed by sonication, clarified by centrifugation at 20,000 g for 30 minutes, then bound to 5 mL of Ni-NTA resin (Thermo Scientific 88222) for 1 hour at 4°C. The resin was washed with 100 mL of His binding buffer, then the protein was eluted in 20 mL of His elution buffer (20 mM HEPES pH 7.5, 150 mM KCl, 2.5 mM MgCl_2_, 250 mM imidazole, and 1 mM BME). The protein was concentrated in a spin concentrator, then loaded onto a Superdex 200 16/60 column (GE Healthcare) equilibrated in buffer (20 mM HEPES pH 7.5, 150 mM KCl, 2.5 mM MgCl_2_, 10 % glycerol, and 0.5 mM TCEP). Fractions corresponding to the monomeric protein were pooled together, concentrated, and aliquoted for storage at −80 °C.

### Purification of ClpX and ClpP

The purification of ClpXΔN and ClpP were done as previously described (Martin et al., 2005). A plasmid encoding a linked trimer of ClpXΔN with an N-terminal 6xHis affinity tag was transformed into *E. coli* BL21(DE3) and grown in 1 L of TB until OD_600_ of 0.6. Protein production was induced with 0.5 mM IPTG. Cells were harvested after 4 hours at 37°C and resuspended in His binding buffer (20 mM HEPES pH 7.5, 100 mM KCl, 400 mM NaCl, 20 mM imidazole, 10% glycerol, and 1 mM BME), supplemented with protease inhibitors and Pierce Universal Nuclease (Thermo Scientific PI88702). Cells were lysed by sonication, clarified by centrifugation at 20,000 g for 30 minutes, then bound to 5 mL of Ni-NTA resin (Thermo Scientific 88222) for 1 hour at 4_°_C. The resin was washed with 100 mL of His binding buffer, then the protein was eluted in 20 mL of His elution buffer (20 mM HEPES pH 7.5, 100 mM KCl, 400 mM NaCl, 250 mM imidazole, 10% glycerol, and 1 mM BME). The protein was concentrated in a spin concentrator, then loaded onto a Superdex 200 16/60 column (GE Healthcare) equilibrated in buffer (20 mM HEPES pH 7.5, 300 mM KCl, 0.1 mM EDTA, 10 % glycerol and 1 mM DTT). Fractions corresponding to the monomeric protein were pooled together, concentrated and aliquoted for storage at −80°C. Protein concentration was determined by measuring A_280_.

A plasmid encoding ClpP with a C-terminal 6xHis affinity tag was transformed into *E. coli* BL21(DE3), grown in 1 L of TB until OD_600_ of 0.6. Protein production was induced with 0.5 mM IPTG, and cells were harvested after 4 hours at 37°C, and resuspended in His binding buffer (20 mM HEPES pH 7.5, 100 mM KCl, 400 mM NaCl, 20 mM imidazole, 10% glycerol, and 1 mM BME), supplemented with Pierce Universal Nuclease (Thermo Scientific PI88702). Protease inhibitors were omitted. Cells were lysed by sonication, clarified by centrifugation at 20,000 g for 30 minutes, then bound to 5 mL of Ni-NTA resin (Thermo Scientific 88222) for 1 hour at 4°C. The resin was washed with 100 mL of His binding buffer, then the protein was eluted in 20 mL of His elution buffer (20 mM HEPES pH 7.5, 100 mM KCl, 400 mM NaCl, 250 mM imidazole, 10% glycerol, and 1 mM BME). The protein was then bound to a 5 mL HiTrap MonoQ column equilibrated in low salt buffer (50 mM Tris-HCL pH 8.0, 50 mM KCl, 10 mM MgCl_2_, 0.1 mM EDTA, 10% glycerol, and 1 mM BME), washed with 20 mL of low salt buffer and then eluted using a 100 mL gradient between low salt buffer and high salt buffer (50 mM Tris-HCL pH 8.0, 200 mM KCl, 10 mM MgCl_2_, 0.1 mM EDTA, 10% glycerol, and 1 mM BME). The protein was concentrated in a spin concentrator, then loaded onto a Superdex 200 16/60 column (GE Healthcare) equilibrated in buffer (20 mM HEPES pH 7.5, 200 mM KCl, 0.1 mM EDTA, 10 mM MgCl_2_, 10 % glycerol and 1 mM DTT). Fractions corresponding to the monomeric protein were pooled together, concentrated and aliquoted for storage at −80°C. Protein concentration was determined by measuring A_280_.

### Purification of remaining recombinant proteins

The rest of the recombinant proteins used in this paper were expressed with an N-terminal 6xHis-SUMO tag in *E. coli* strain BL21(DE3) and purified as described for Pab1, but using anion exchange instead of Heparin column.

### Fluorescein labeling of Pab1

Fluorescein labeling of Pab1 termini was done using sortase catalyzed ligation of labeled peptides. For N-terminal labeling, 6xHis-TEV-GGG-Pab1 was purified using the same protocol described for wild-type Pab1 above. The labeling was done in SEC buffer (20 mM HEPES 7.3, 150 mM KCl, and 2.5 mM MgCl_2_) as a 500 *µ*L reaction with 100 *µ*M Pab1, 20 *µ*M wild-type sortase A, 0.5 mM TCEP, 10 mM CaCl_2_ and 0.5 mM 5-FAM-HHHHHHLPETGG peptide (Biomatik). After an hour incubation at room temperature, labeled protein was separated from free peptide on a 5 mL HiTrap Desalting column equilibrated in aggregation buffer (20 mM HEPES pH 6.8, 150 mM KCl, 2.5 mM MgCl_2_, and 1 mM DTT). The C-terminally labeled Pab1 was prepared similarly, but the labeling reaction was done with the following condition: 100 *µ*M Pab1, 20*µ*M heptamutant sortase A, 0.5 mM TCEP, and 0.5 mM GGGK(FAM)AANDENYALAA or GGGK(FAM) peptide (Biomatik).

### In vivo Total/Soluble/Pellet (TSP) assay

Yeast cells were diluted to OD_600_ of approximately 0.001 in 250 mL YPD and incubated at 30°C until OD_600_ reached between 0.3 and 0.4. 30 mL of the cell culture was harvested as the pre-shock sample by spinning in a 50 mL conical tube at 3,000 g for 5 minutes at room temperature (RT). The cell pellet was resuspended in 1 mL of cold soluble protein buffer (SPB; 20 mM HEPES pH7.3, 120 mM KCl, 2 mM EDTA, 0.2 mM DTT, 1:100 PMSF, and 1:100 protease inhibitors cocktail IV (MilliporeSigma 539136)), transferred to a pre-chilled 1.5 mL microcentrifuge tube, and centrifuged again at 5,000 g for 30 seconds at RT. Supernatant was removed and the pellet was resuspended in 170 *µ*L SPB. Two 100 *µ*L aliquots from the resuspended sample were snap-frozen in liquid nitrogen. The remaining cell culture was vacumm filtered and the cell pellet was transferred to a 50 mL conical tube. The conical tube was placed in a 42°C water bath to heat shock the cells. After 20 minutes, the cell pellet was resuspended in 220 mL of pre-warmed 30°C YPD media. 30 mL of this culture was harvested as the heat shock sample and processed as described earlier. The remaining cell culture was transferred to a 1 L flask and the cells were recovered in a 30°C water bath. Recovery samples were collected at different time points and processed in the same way as the pre-shock sample. Cells were lysed by cryomilling and fractionated as described in Wallace et al. (2015), with minor modifications on spin conditions. Briefly, cell lysates were cleared at 3,000 g for 30 seconds. 150 *µ*L of the cleared lysate was transferred to a new 1.5 mL tube. To remove RNA, RNase I_f_ (NEB M0243S) was added to the final concentration of 0.3 units/*µ*L and the sample was incubated at RT for 30 minutes. 50 *µ*L of the sample was transferred to a new tube, mixed with Total protein buffer (TPB; 20 mM HEPES pH 7.3, 150 mM NaCl, 5 mM EDTA, 3% SDS, 1:100 PMSF, 2 mM DTT, and 1:1000 protease inhibitors IV (MilliporeSigma 539136)), and spun at 6,000 g for 1 minute to collect Total protein sample. Pellet samples were collected by spinning the remaining 100 *µ*L sample at 8,000 g for 5 minutes (P_8_) and at 100,000 g for 20 minutes (P_100_) at 4°C, and resuspending the pellet in Insoluble protein buffer (IPB; TPB with 8 M urea). Supernatant from this last spin was collected as the Soluble protein sample. Pab1 in each sample were visualized by SDS-PAGE and western blot as described in (Wallace et al., 2015) using mouse monoclonal anti-Pab1p antibody (EnCor Biotechnology MCA-1G1) and Image Lab software (Bio-Rad).

### In vitro Total/Soluble/Pellet (TSP) assay

Reaction mixture containing Pab1 condensates and molecular chaperones were prepared and incubated at 30°C for an hour either in the absence or presence of 5 mM ATP. The reaction mixture (Table 2) were centrifuged at 100,000 g for 20 minutes at 4°C. Supernatant was collected as the soluble fraction sample. Buffer (20 mM HEPES pH 7.3, 150 mM KCl, 2.5 mM MgCl_2_, 0.01 % Triton X-100, 0.5 mg/mL BSA, and 1 mM DTT) was added to the pellet and the sample was centrifuged again under the same condition. After removing the supernatant, the pellet was directly resuspended in 1x Laemmli buffer. Pab1 in each sample was visualized by SDS-PAGE and western blot as described in (Wallace et al., 2015) using Image Lab software (Bio-Rad).

**Table 2.**
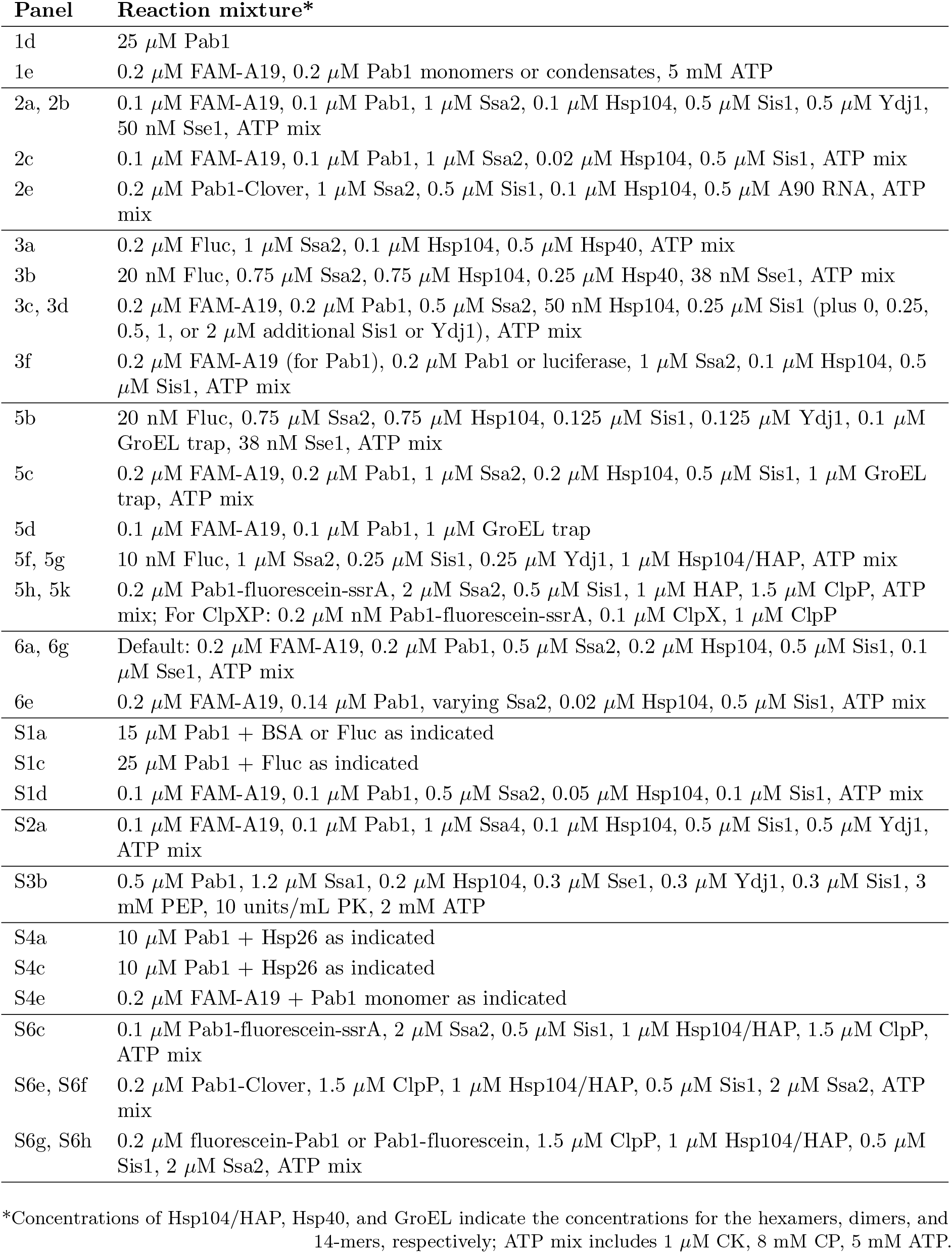
Reaction conditions for experiments shown in the figures.

### Preparation of Pab1 condensates

Pab1 monomers were buffer exchanged into aggregation buffer (20 mM HEPES pH 6.8, 150 mM KCl, 2.5 mM MgCl_2_, and 1 mM DTT) and diluted to make a 500 *µ*L sample of 25 *µ*M Pab1. The sample was sometimes supplemented with 100-fold lower firefly luciferase to increase yield. The sample was incubated in a 42°C water bath for 20 minutes at pH 6.8. Under this heat shock condition, the solution remained clear and minimal protein pelleting was observed after a 3 minute centrifugation at 8,000 g. N-terminally fluorescein labeled Pab1 aggregates were prepared at 39°C. The supernatant was loaded onto a Superose 6 10/300 GL column (GE Healthcare) equilibrated with SEC/Storage buffer (20 mM HEPES pH 7.3, 150 mM KCl, 2.5 mM MgCl_2_, and 1 mM DTT). Void fractions containing small Pab1 assemblies (*>*5,000 kDa) were collected. About half of the loaded protein eluted in the void volume while the remaining protein eluted as monomers. Concentration of Pab1 condensates in each void fraction was measured using Bradford assay and/or SDS-PAGE with Pab1 standards of known concentrations. Protein aliquots were snap-frozen in liquid nitrogen and stored at −80°C.

### Monitoring Pab1 dispersal using fluorescence anisotropy

Because Pab1 binds 12-mer poly(A) with full affinity and has a binding footprint of roughly 25 nucleotides (Sachs et al., 1987), we used 19-mer poly(A) RNA to get 1:1 binding of Pab1 to RNA. Pab1 condensates, molecular chaperones, and 5’ labeled A19 RNA (FAM-A19 or Atto550-A19) were diluted to desired concentrations in disaggregation buffer (20 mM HEPES pH 7.3, 150 mM KCl, 2.5 mM MgCl_2_, 0.5 mg/mL BSA, 0.01 % Triton X-100, and 1 mM DTT). The reaction mixture (Table 2) was supplemented with 5 mM ATP, and 8 mM creatine phosphate (CP) and 1 *µ*M creatine kinase (CK) for ATP regeneration. The reaction mixture was also supplemented with 2 % SUPERase RNase Inhibitor (Thermo Fisher AM2694) to prevent RNA degradation. The final reaction volume was 15 *µ*L. Calibration samples were prepared by adding known concentrations of monomeric Pab1 to a fixed concentration of FAM-A19 used in the reaction samples, typically 200 nM.

A new calibration curve was measured each time an experiment was performed. The reaction mixtures were added to non-binding surface 384-well plate (Corning CLS3575). The plate wells were sealed with a plate sealer (Thermo Fisher 235307) to prevent liquid evaporation. Fluorescence anisotropy was measured every 20 second in Spark microplate reader (TECAN) using excitation/emission wavelengths of 485 nm/535 nm, each with a bandwidth of 20 nm, at 30°C. Disaggregation buffer was used as blank. G factor was calibrated with a solution of free 6-iodoacetamidofluorescein to produce a fluorescence polarization reading of 20 mP.

### Anisotropy data fitting and analysis

The reaction data were fitted with the following logistic equation:

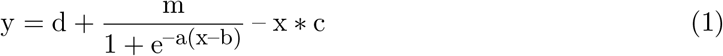

where d, m, a, b, and c are fitting parameters. The negative linear term accounts for the chaperone concentration-dependent signal decay, which comes from RNA-degrading contaminants copurified with chaperones.

To extract maximal rate of Pab1 dispersal, the fluorescence anisotropy values were first converted to the concentration of RNA-binding competent monomeric Pab1 using the calibration curve. Fluorescence anisotropy data of the calibration samples were fitted with the following equation:

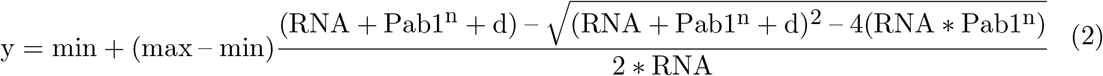

Min and max refer to the fluorescence anisotropy values of the calibration samples with no or saturating amount of monomeric Pab1, respectively. The values of d and n extracted from the calibration fit were used to convert fluorescence anisotropy values in the reaction samples to concentrations of Pab1 with this rearranged equation (2):

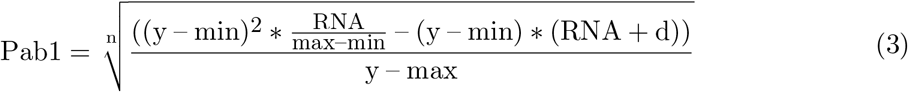

The converted data were fitted again with equation (1). Maximal rate of dispersal was calculated by computing the extracted fit parameters to the derivative of equation (1) when x = b:

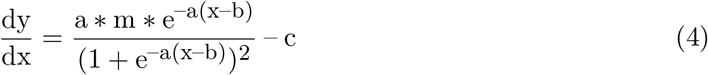

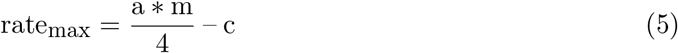

Note that, because Pab1 condensates exhibit substantially reduced but non-zero binding to FAM-A19 in a concentration-dependent manner, the maximal rate of dispersal quantified from fluorescence anisotropy data may be systematically underestimating the true rate of dispersal.

To convert the y-axis from Pab1 concentration to fraction restored Pab1, we first subtracted background signal using the negative control data (no ATP). Background-subtracted data were divided by the total concentration of Pab1, which was approximated by taking the mean of highest 50-100 data points, i.e., data points in the plateau region of the positive control. Total Pab1 concentration in the reaction had to be approximated this way for more accurate quantification because we noticed that Pab1 condensates adhere to plastic, causing loss of about 30-50% substrate during transfer. To compensate for this, 1.5 to 2-fold excess Pab1 was added to aim for, e.g., the final concentration of 0.2 *µ*M Pab1 in the reaction. The same total Pab1 concentration was used to normalize reactions prepared from the same master mix. The presence of 5 mM ATP slightly lowered the fluorescence anisotropy baseline compared to the no ATP control, and this led to negative starting values for all ATP-containing reactions after background subtraction; All traces were shifted upward by the same amount to make the positive control reactions to start around the value of zero.

The rate data in Figure 6A were fitted with a logistic equation:

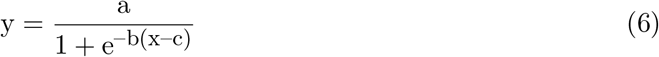

with a, b, and c as fitting parameters. We used the total Pab1 concentration to calculate the ratio of chaperone to substrate.

### Fluorescence-detection size-exclusion chromatography (FSEC)

Small Pab1-Clover condensates were prepared and mixed with chaperones, ATP, ATP regeneration system, and SUPERase RNase Inhibitor as described above for wild type Pab1 (Table 2). The reaction samples (120 *µ*L per run) were incubated at 30°C for an hour. 120 *µ*L of the sample was loaded to a Superdex 200 10/300 GL column (GE Healthcare) equilibrated with filtered running buffer (20 mM HEPES pH 7.3, 150 mM KCl, 2.5 mM MgCl_2_, and 1 mM DTT) using Akta system. Fluorescence was measured by a fluorescence detector (JASCO FP-2020 Plus) connected to the Akta system.

### Luciferase reactivation assay

Recombinant firefly luciferase was aggregated by incubating 2 *µ*M of luciferase with 10 *µ*M Hsp26 in aggregation buffer (20 mM HEPES pH 6.8, 150 mM KCl, 2.5 mM MgCl_2_, and 1 mM DTT) at 42°C for 20 minutes. After cooling on ice for 2 minutes, the aggregates were diluted to 0.2 *µ*M in the disaggregation buffer (20 mM HEPES pH 7.3, 150 mM KCl, 2.5 mM MgCl_2_, 0.5 mg/mL BSA, 0.01 % Triton X-100, and 1 mM DTT) supplemented with 5 mM ATP, 8 mM CP, 1 *µ*M CK, and specified concentrations of chaperones (Table 2). The mixed sample was incubated at 30°C. At each time point, 2 *µ*L of the reaction sample was added to 18 *µ*L of luciferin mix (Promega E1500), and luminescence was measured using Spark microplate reader (TECAN) with integration time of 1 second. Luminescence of 0.2 *µ*M native luciferase supplemented with 1 *µ*M Hsp26 in disaggregation buffer was used to compute reactivation yield.

For faster disaggregation using excess molecular chaperones (Figure 3B), 33.8 nM luciferase was heat shocked in the presence of 169 nM Hsp26 at 42°C for 20 minutes in low-salt aggregation buffer (25 mM HEPES pH 7.3, 50 mM KCl, 0.1 mM EDTA, and 1 mM DTT). Luciferase aggregates were diluted to the final concentration of 20 nM in the disaggregation buffer containing chaperones (Table 2). Luminescence was measured as described above.

For disaggregation of chemically aggregated luciferase using HAP/ClpP system (Figure 5H and 5I), luciferase was diluted to 5 *µ*M in low-salt urea buffer (25 mM HEPES pH 7.3, 50 mM KCl, 0.1 mM EDTA, 1 mM DTT, and 8 M urea) and incubated at 30°C for 30 minutes. Aggregation was induced by diluting luciferase 100-fold into the disaggregation buffer containing chaperones and HAP/ClpP (Table 2). Western blot samples were collected at the specified time points and stained using luciferase antibody (MilliporeSigma L0159). Blots were visualized using Odyssey CLx (LI-COR). Luminescence was measured as described above using mock-treated luciferase as normalization control.

### Dynamic light scattering (DLS)

DLS measurements were performed using DynaPro NanoStar (Wyatt). Sample acquisition was done as described in (Riback et al., 2017). All experiments, unless noted otherwise, were performed with 10 *µ*M Pab1 in filtered DLS buffer (20 mM HEPES pH 6.8, 150 mM KCl, 2.5 mM MgCl_2_, and 1 mM DTT). All protein samples used for DLS experiments were dialyzed against DLS buffer overnight at 4°C and cleared of aggregates by spinning at 20,000 g for 20 minutes.

### GroEL trap assay

Pab1 dispersal and luciferase reactivation assays were done as described above, but in the presence of 5-fold excess GroEL trap (Table 2). For refolding experiment, 5 *µ*M Pab1 was denatured in 8 M urea buffer (20 mM HEPES pH 6.8, 150 mM KCl, 2.5 mM MgCl_2_, 1 mM DTT, and 8 M urea) and incubated at 30°C for 30 minutes. Pab1 was first diluted to 0.5 *µ*M in refolding buffer (20 mM HEPES pH 7.3, 150 mM KCl, 2.5 mM MgCl_2_, and 1 mM DTT) containing no or 10-fold excess GroEL trap, and then to 0.1 *µ*M in the same respective refolding buffer with 0.1 *µ*M FAM-A19. Pab1’s RNA-binding activity was measured by fluorescence anisotropy.

### Gel analysis of HAP/ClpP degradation

Digestion by the HAP/ClpP system was done in disaggregation buffer (20 mM HEPES pH 7.3, 150 mM KCl, 2.5 mM MgCl_2_, 0.5 mg/mL BSA, 0.01 % Triton X-100, and 1 mM DTT) with 0.2 *µ*M Pab1 condensate or monomer, 1.5 *µ*M ClpP, 1 *µ*M Hsp104 (WT or HAP), 0.5 *µ*M Sis1, 1-2 *µ*M Ssa2, 5 mM ATP, and 8 mM CP and 1 *µ*M CK for ATP regeneration (Table 2). Reactions were run for 1 hour at 30°C, then quenched with Laemmli buffer and run on a Bio-rad TGX 4-20% SDS-PAGE gel. Fluorescent gels were imaged using a ChemiDoc (Bio-rad) and westerns were performed using a 1:5000 dilution of Rabbit anti-GFP antibody (Life A11122) and a 1:20,000 dilution of secondary (Donkey anti-Rabbit, LiCor 925-32213). Blots were visualized using Odyssey CLx (LI-COR). The ClpXP digestion reaction was done for 30 minutes at 30°C with 0.2 *µ*M Pab1-FAM monomer, 0.1 *µ*M ClpX and 1 *µ*M ClpP in disaggregation buffer.

### Disaggregation data from the literature

The following data from 18 different studies (Nillegoda et al., 2017; Doyle et al., 2015; Rampelt et al., 2012; Yu et al., 2015; Żwirowski et al., 2017; Martín et al., 2014; Kłosowska et al., 2016; Reidy et al., 2014; Rosenzweig et al., 2013; DeSantis et al., 2012; Shorter, 2011; Haslberger et al., 2008; Glover and Lindquist, 1998; Ratajczak et al., 2009; Cashikar et al., 2005; Goloubinoff et al., 1999; Sielaff and Tsai, 2010; Duennwald et al., 2012) were compiled for comparison: 1) substrate identity; 2) concentrations of substrate and molecular chaperones used; 3) maximum yield observed within the experimental time; 4) the names of molecular chaperones; and 5) reference to the source paper with DOI. Only the results from *in vitro* experiments were recorded. Studies reporting fold-change relative to negative control were omitted because yield cannot be determined from the given information for comparison. For a study which reports multiple disaggregation results with the same substrate, the concentrations of substrate and chaperones which give the highest maximal yield were recorded.

### Modeling and simulation

In our model, free substrates exist in one of the following states: folded (S_f_), unfolded (S_u_), misfolded (S_m_), and aggregated (S_a_). Free Hsp70 exists either in an ATP-bound state (Hsp70_ATP_) or an ADP-bound state (Hsp70_ATP_), and each state can bind certain free substrates to form a complex (e.g., Hsp70_ATP_:S_a_). ATP hydolysis of Hsp70_ATP_ in Hsp70_ATP_:S_m_ complex results in substrate unfolding, a step we call “priming”. For an aggregated substate, the same sequence of events do not result in any state transition but instead primes the complex (Hsp70_ATP_:S_ap_) for interaction with Hsp104. In the cooperative model, a second Hsp70_ATP_ can bind Hsp70_ADP_:S_ap_ complex to form a ternary complex (Hsp70_ATP,ADP_:S_ap_) and only the ternary complex with both Hsp70s in the ADP-bound state (Hsp70_ADP,ADP_:S_ap_) can engage with Hsp104 for disaggregation. For simplicity, we did not allow unfolding of a natively folded substrate.

The time evolution of the concentrations of all distinct species in the cooperative model was described using the following ODEs:

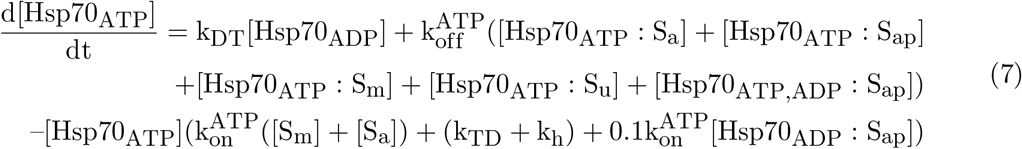

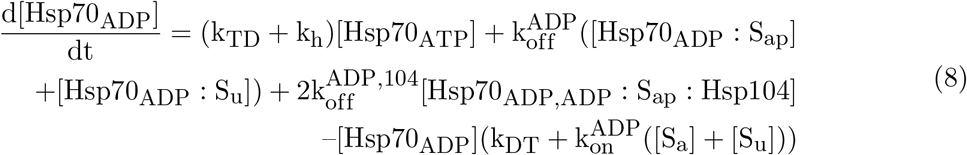

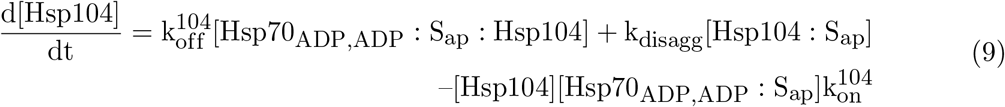

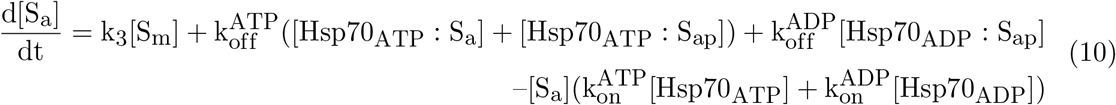

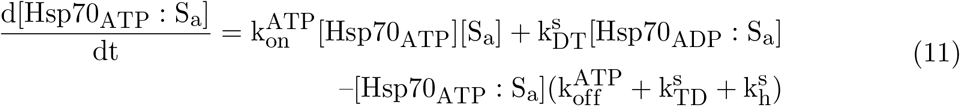

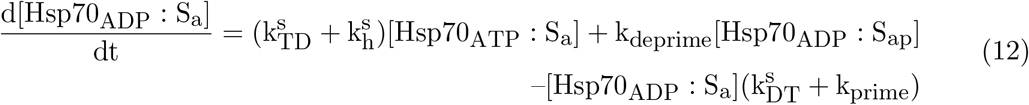

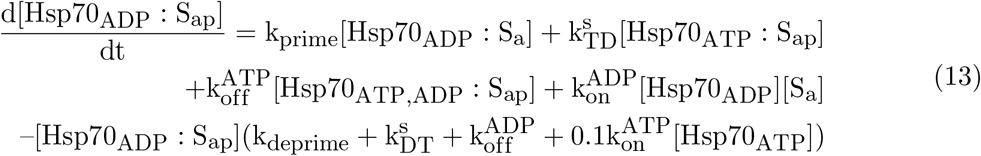

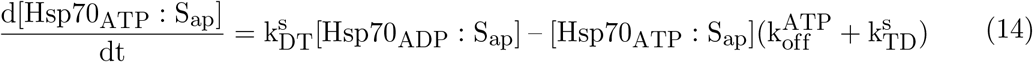

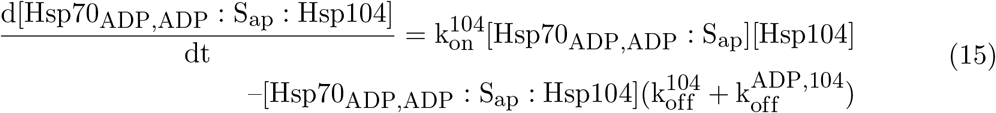

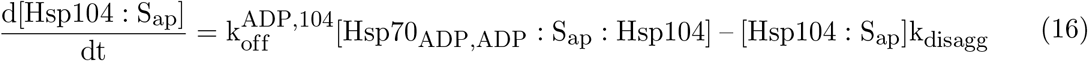

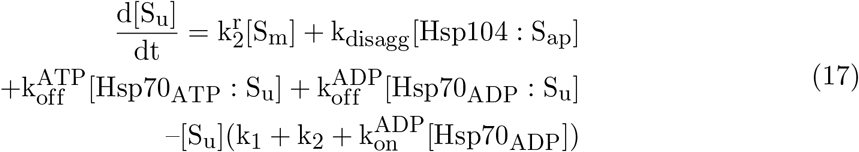

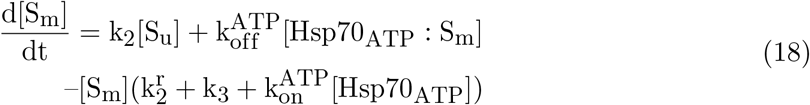

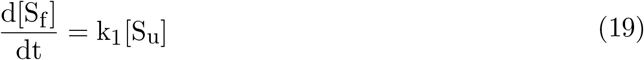

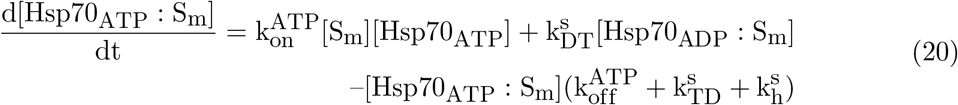

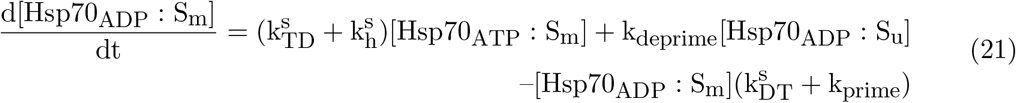

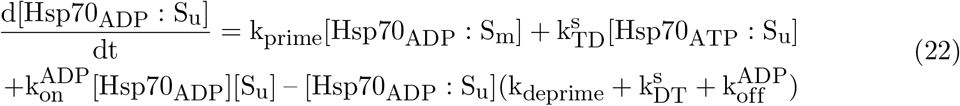

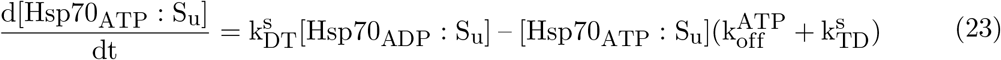

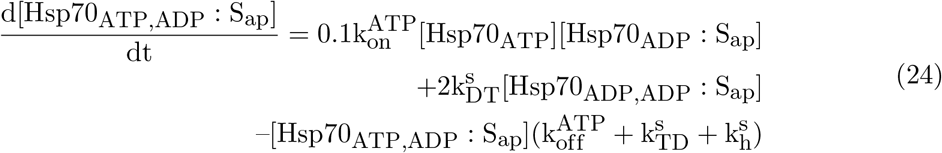

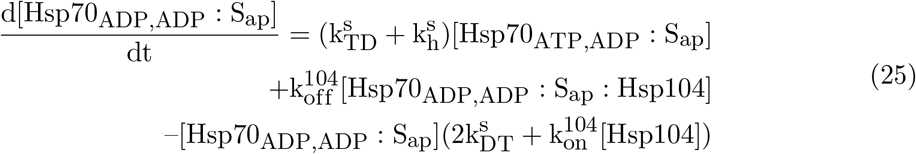

The parameter values used for the simulation are listed in Table 1. Note that the current model lacks spatial information; an extension of this model allowing spatial segregation of aggregated substrates would allow investigating the effect of aggregate shape and/or size on the efficiency of the disaggregation system.

## Supplemental Figures

**Figure S1.**
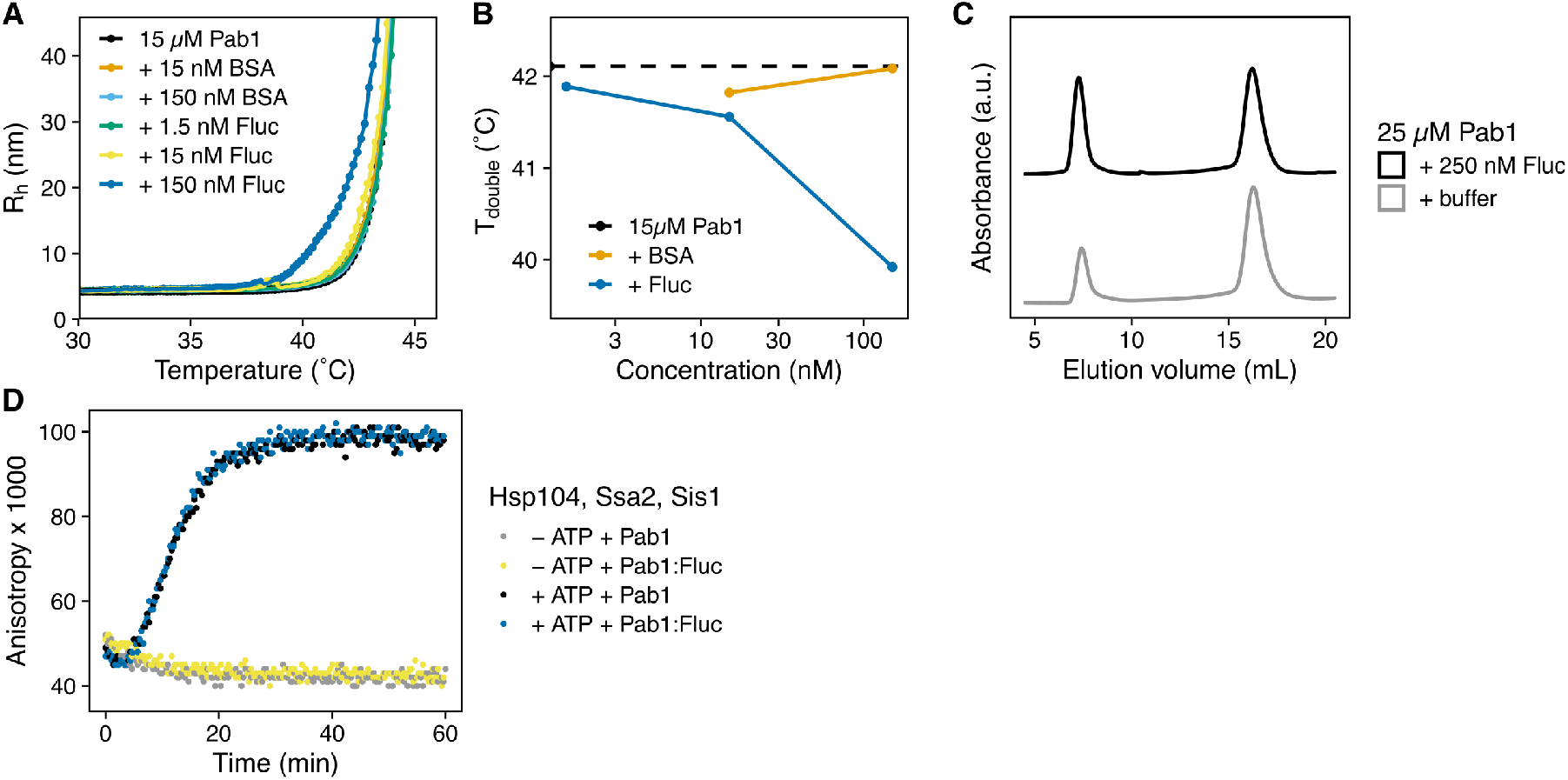
Misfolded protein can nucleate Pab1 condensation; Related to Figure 1. (A) DLS temperature ramp experiment with 15 *µ*M Pab1 in the presence of increasing concentrations of firefly luciferase (Fluc) or BSA. (B) T_double_, temperature at which the baseline R_h_ doubles, for the DLS temperature ramp experiment shown in (A) is plotted against the concentrations of the additives (either BSA or luciferase). Dashed line indicates T_double_ for Pab1 in the absence of any additives. (C) Representative SEC trace of Pab1 heat shocked in the absence or presence of 100-fold lower luciferase. Pab1 condensates and monomers elute around 7.5 mL and 16 mL, respectively. (D) Dispersal of Pab1 condensates formed in the presence or absence of 100-fold less luciferase. Condensation in the presence of 100-fold lower luciferase does not affect subsequent condensate dispersal by chaperones.

**Figure S2.**
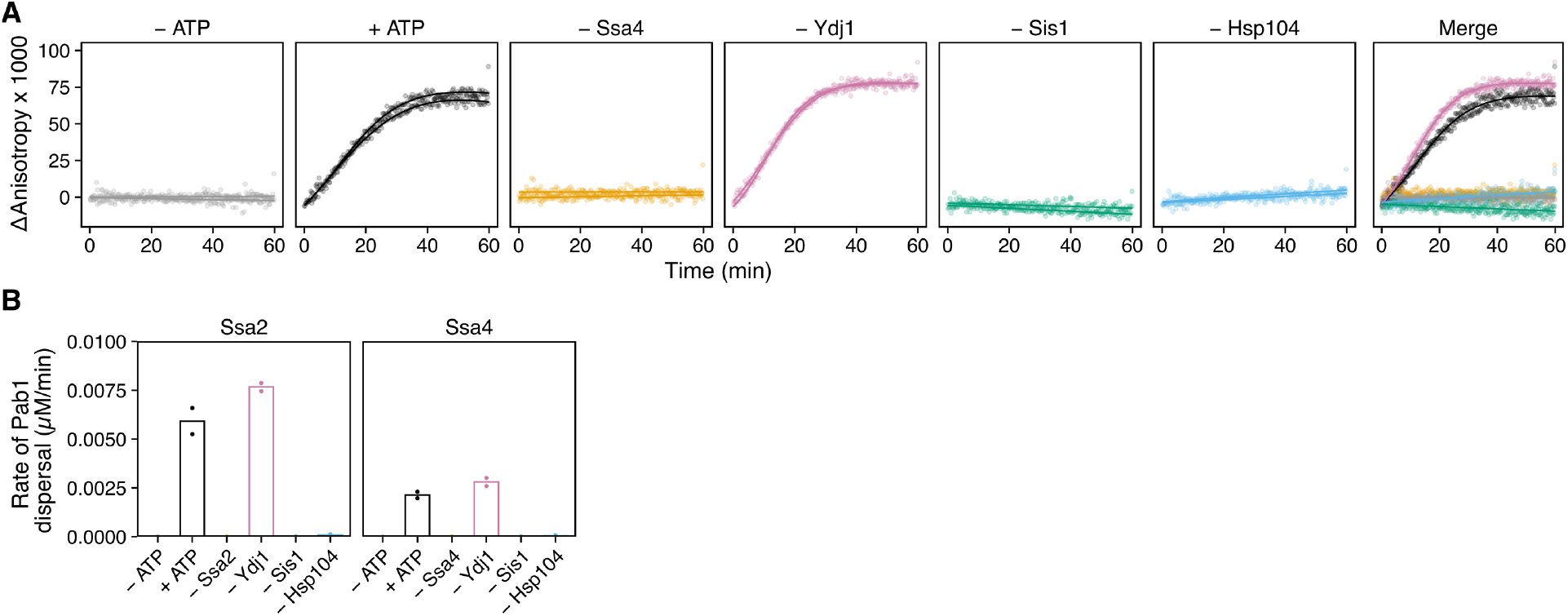
Ssa2 can be replaced by its heat-inducible paralog Ssa4 for Pab1 dispersal; Related to Figure 2. (A) Time-resolved fluorescence anisotropy of A19 in the presence of Pab1 condensates and molecular chaperones. All chaperones shown in Figure 2A, except Sse1, were included in the experiments. Ssa4 was added instead of Ssa2. Fitted data points from two independent experiments are shown. Merged data points and a solid line fitted to the average of each experimental condition is shown on the right. (B) Maximal rate of dispersal quantified from the dropout experiments in the absence of Sse1.

**Figure S3.**
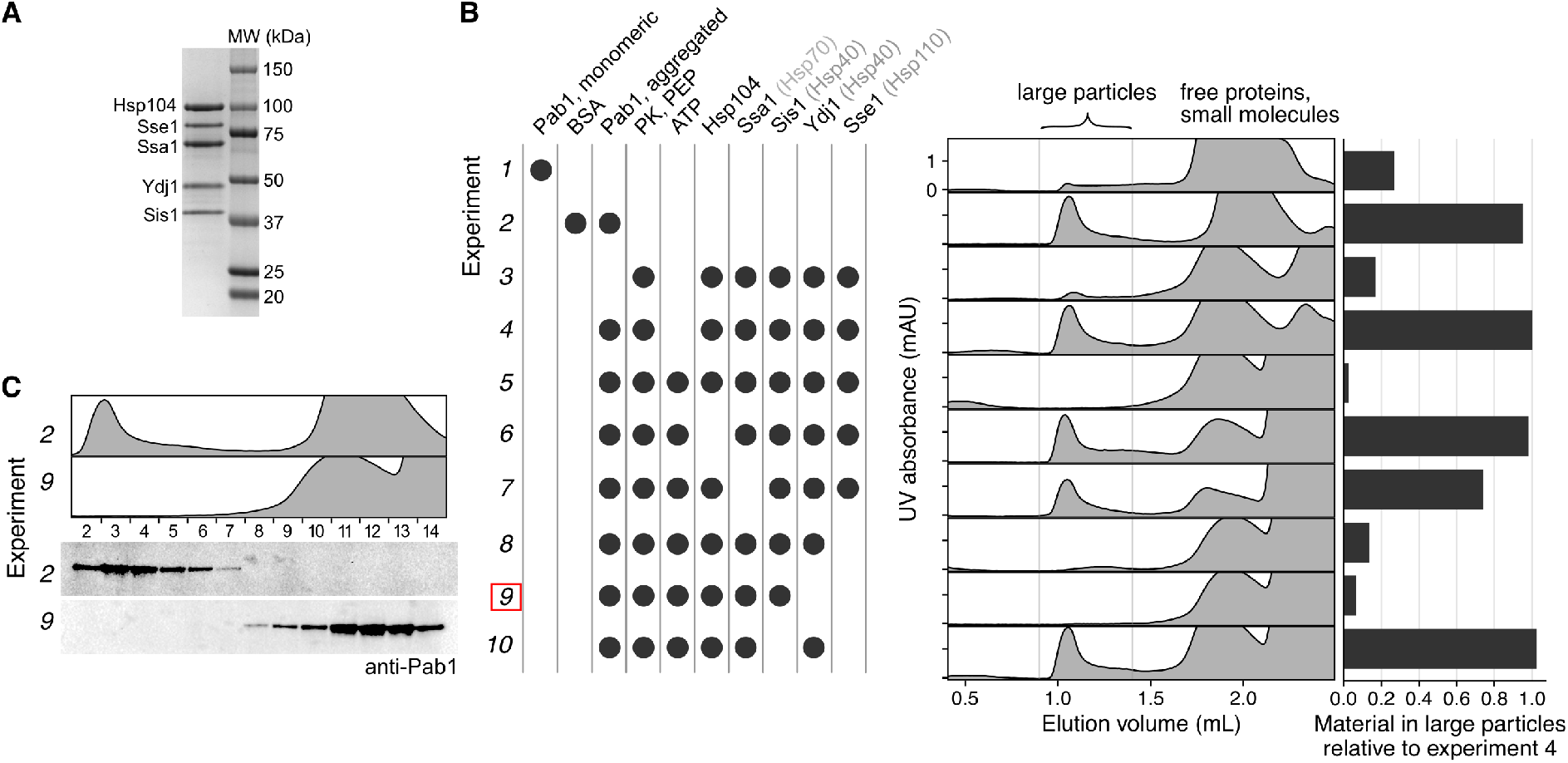
Hsp104, Hsp70, and Sis1 are necessary and sufficient for Pab1 dispersal; Related to Figure 2. (A) Purification of recombinant chaperones. (B) Dropout experiments to determine the components necessary for dispersal of Pab1 condensates. Pab1 dispersal reaction samples were examined by Superose 6 Precision Column. Red box highlights the minimal set composed of Hsp104, Ssa1, and Sis1. PK and PEP stand for pyruvate kinase and phosphoenolpyruvate, respectively. (C) Verification of Pab1 dispersal by western blot.

**Figure S4.**
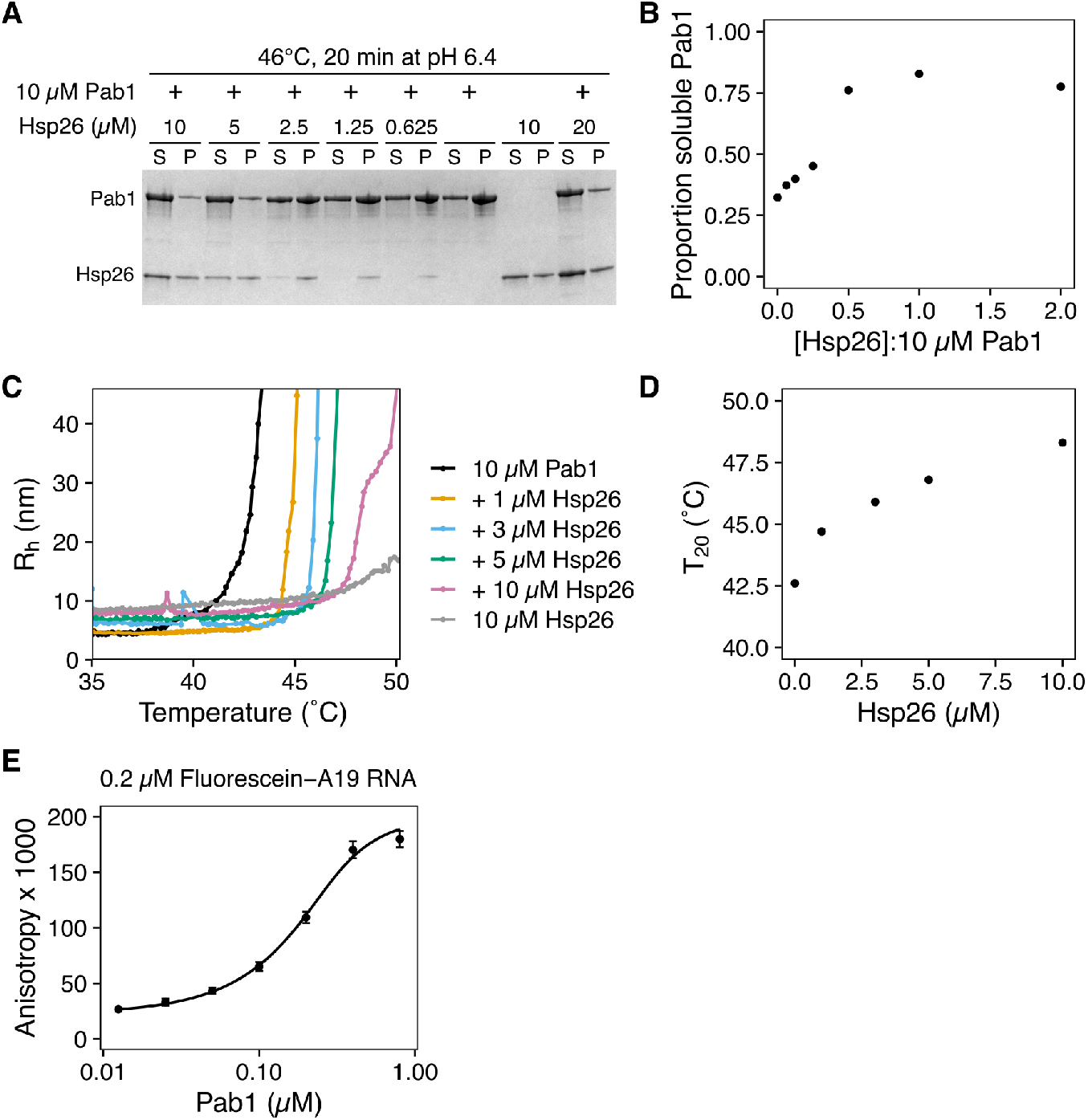
Hsp26 suppresses Pab1 condensation; Related to Figure 3. (A) DLS temperature ramp experiment with 10 *µ*M Pab1 with increasing concentrations of Hsp26 at pH 6.4. Note that the presence of Hsp26, which forms high molecular weight oligomers, shifts the DLS baseline upward. (B) Temperature at which Pab1’s hydrodynamic radius crosses 20 nm in (A) is plotted against Hsp26 concentration. (C) Sedimentation of Pab1 in the absence or presence of increasing concentrations of Hsp26, visualized by Coomassie staining. (D) Quantification of proportion soluble Pab1 in (C) plotted against the ratio of Hsp26 to Pab1. (E) A representative calibration curve showing the increasing fluorescence anisotropy of the labeled A19 RNA as a function of increasing concentration of monomeric Pab1. The calibration curve was used convert the y-axis from fluorescence anisotropy to Pab1 concentration and extract the rate of dispersal. The mean and standard deviation of 8 independent calibration data are shown. Data were fitted with equation 2.

**Figure S5.**
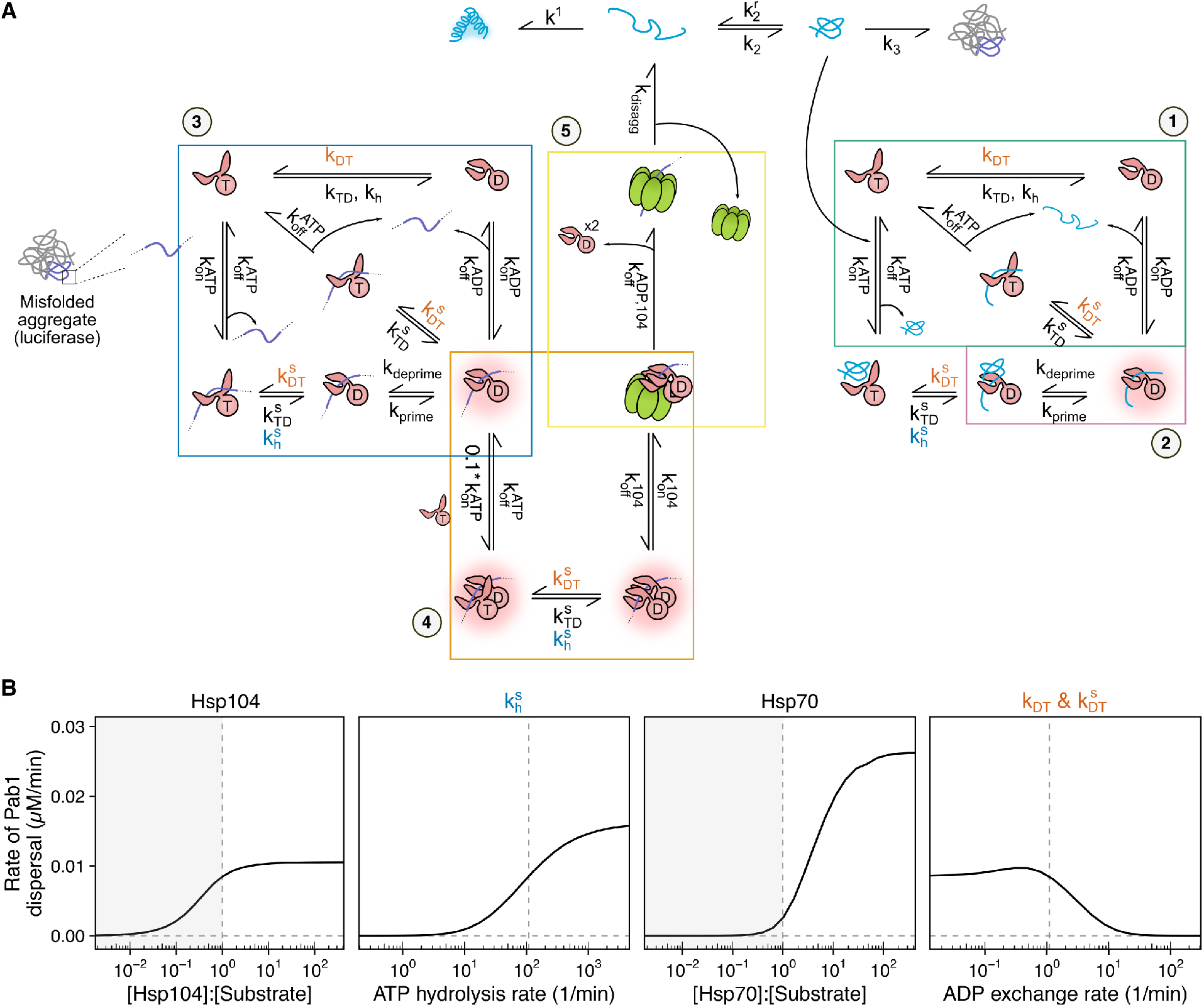
Cooperative model; Related to Figure 4 and Figure 6. (A) (1) Biochemical model of ATP hydrolysis-coupled substrate binding and release from Hsp70 (De Los Rios and Barducci, 2014; Powers et al., 2012; Nguyen et al., 2017; Xu, 2018). Nucleotide exchange from ADP to ATP facilitates substrate release from Hsp70. (2) Hsp70 binding to a single misfolded protein leads to protein unfolding and expansion (Imamoglu et al., 2020; Assenza et al., 2019). We describe this step a “priming” step, and use pink halo to graphically represent the primed species. (3) Nucleotide state-coupled substrate binding of Hsp70 is assumed to be consistent with aggregated substrates. However, unlike with single misfolded proteins, aggregated proteins do not unfold upon Hsp70 binding and need Hsp104. (4) One Hsp70 molecule is insufficient to activate Hsp104 (Seyffer et al., 2012; Carroni et al., 2014). Second Hsp70 binds the substrate with an order of magnitude lower affinity than the first Hsp70 due to the entropic penalty (Wentink et al., 2020). (5) Exact mechanism of substrate handover is unknown, but substrate handover is described as an irreversible step (Powers et al., 2012). (B) Simulation of chaperone titration experiments shown in Figure 6A. Titration of Sis1 and Sse1 were simulated by varying the ATP hydrolysis rate of substrate-bound Hsp70 (kh S) and ADP exchange rate (kDT and kDT S), respectively, from the default level indicated by the dashed line.

**Figure S6.**
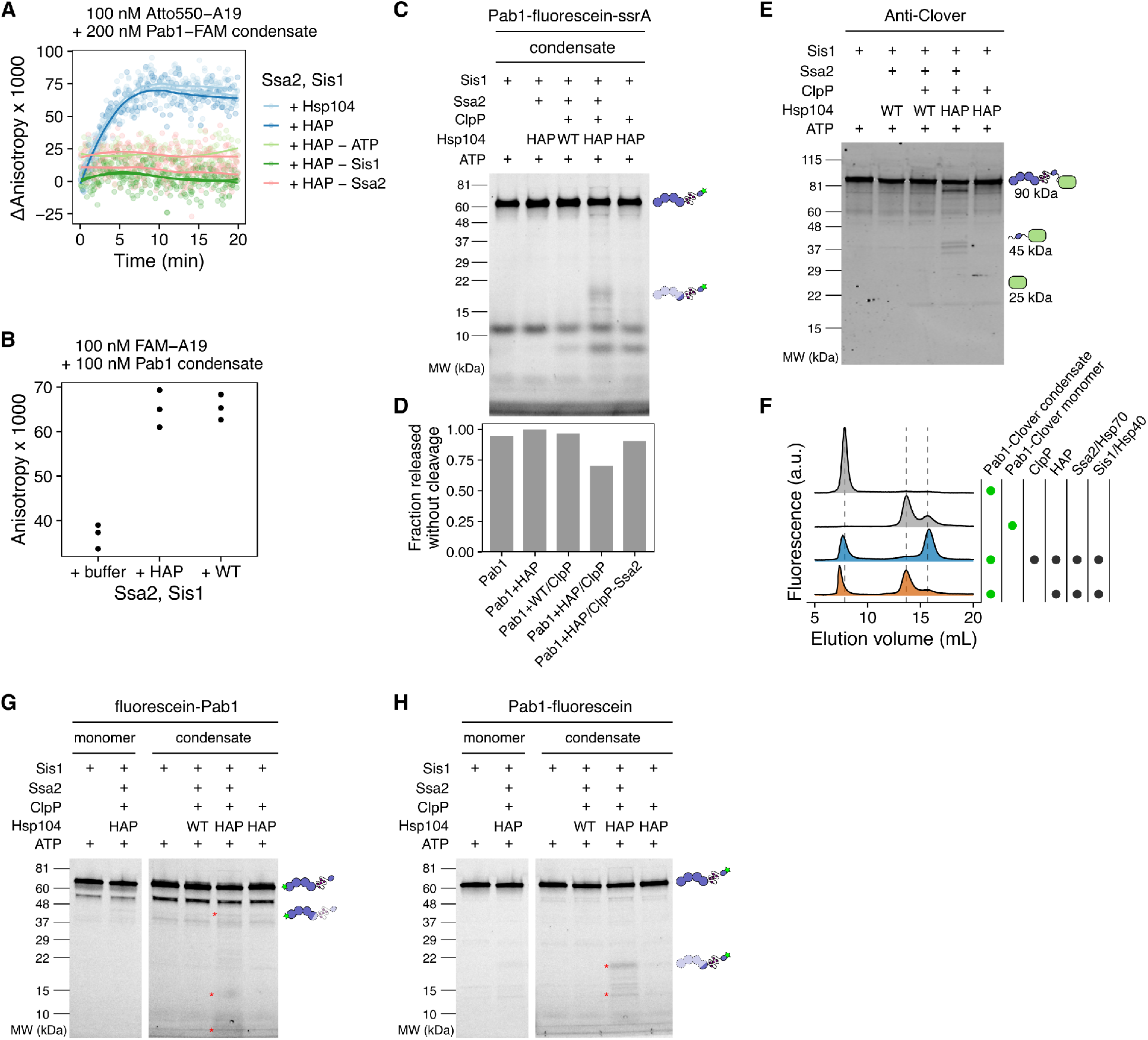
HAP/ClpP-specific cleavage of Pab1 constructs; Related to Figure 5. (A) Condensates of C-terminally fluorescein-labeled Pab1 were dispersed by the indicated components. Fluorescence anisotropy of Atto550-A19 RNA was monitored. (B) End-point measurement of unlabeled Pab1 condensate dispersal. FAM-A19 RNA was added after an hour of reaction incubation to measure fluorescence anisotropy. (C) A replicate of the condensate dispersal experiment shown in Figure 5H.(D) Quantification of full-length Pab1 band intensity in (C) normalized to the HAP control signal. The yield of dispersal was assumed to be the same as seen in the Figure 5K FSEC experiment. (E) Pab1-Clover condensates were incubated with the indicated components and examined by western blot. Schematics of full-length and truncated products, and their corresponding molecular weight are shown. (F) Pab1-Clover condensate was incubated with the indicated components for an hour and examined by FSEC. Dashed lines indicate the elution volume for Pab1-Clover condensates (7.8 mL), Pab1-Clover monomers (13.7 mL), and HAP/ClpP-specific cleavage products (15.7 mL). (G-H) SDS-PAGE gels of N- or C-terminally labeled Pab1 were visualized by detecting fluorescein fluorescence. Asterisks in indicate HAP/ClpP-specific cleavage products.

**Figure S.**
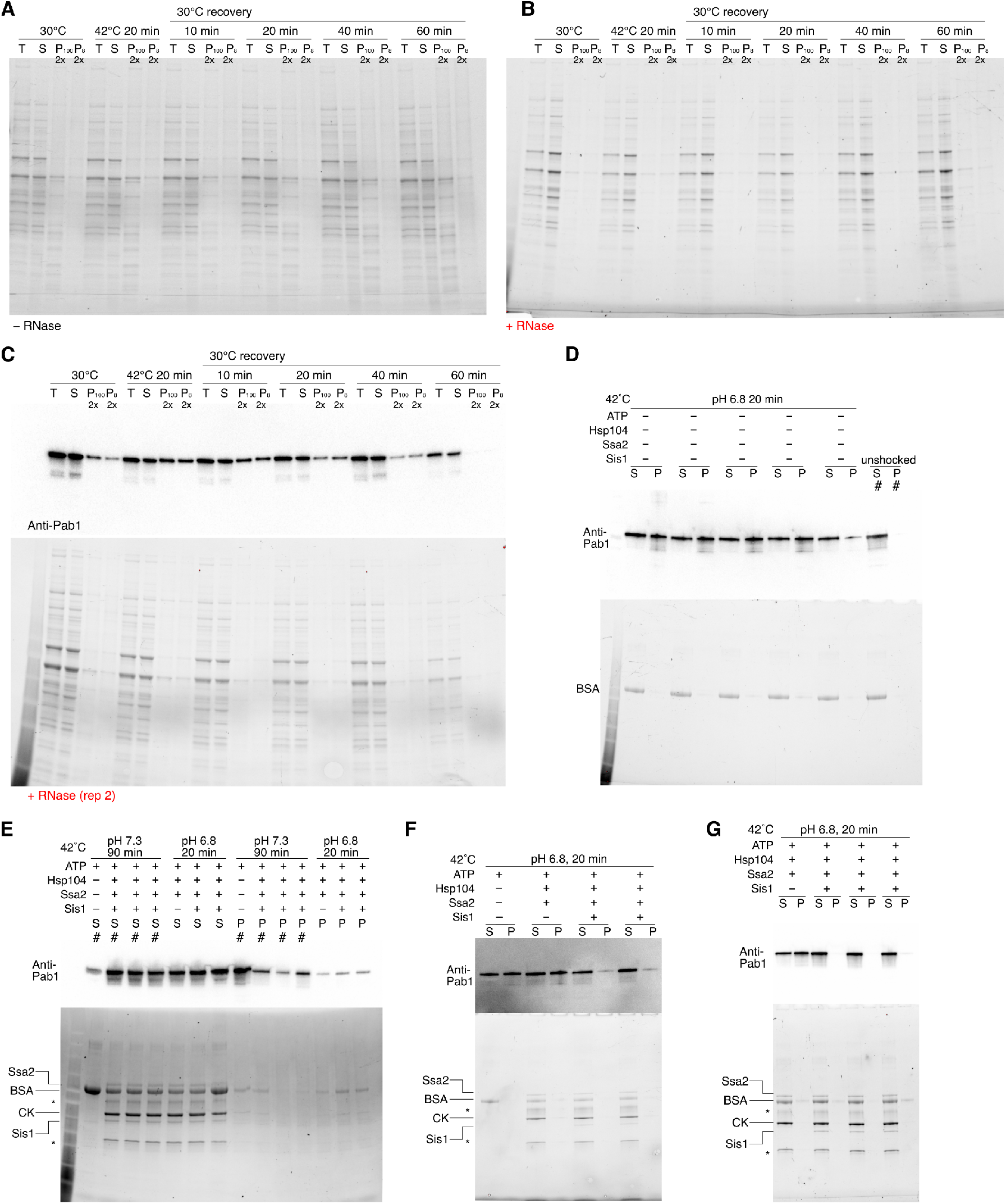
Uncropped stain-free total protein gel and western blot images; Related to Figures 1 and Figure 2. (A and B) SDS-PAGE gel of RNase treated (A) or untreated (B) yeast lysate samples. The corresponding anti-Pab1 western blot images are shown in (Figure 1A). (C) Replicate of (B). (D-G) Western blot (top) and SDS-PAGE images (bottom) of *in vitro* TSP experiments. Cropped images of (G) is shown in (Figure 2C). The quantification results are shown in (Figure 2D). Lanes marked with # were not quantified. CK stands for creatine kinase. Asterisks indicate unknown contaminant.

## Competing Interests

The authors declare no competing interests.

## Acknowledgements

We thank the members of the Drummond lab for helpful comments and discussions. We also thank Tobin Sosnick and Ruofan Chen for helpful comments and discussions, Zachary March for providing the original protocol for Hsp70 purification, Axel Mogk for providing the plasmid for GroEL trap, Andreas Martin for providing the plasmids for ClpX and ClpP, David Pincus and Michael Rust for their feedback on modeling, the Perozo lab for providing the FSEC instrument, and Elena Solomaha at the Biophysics Core for assistance with DLS experiments. Research reported in this publication was supported by the National Institute of General Medical Sciences and the National Institute of Environmental Health Sciences of the National Institutes of Health (NIH) through awards to HY (award numbers T32GM007183 and F31ES030697). JAMB acknowledges fellowship support from the Helen Hay Whitney Foundation. DAD acknowledges support from the NIH (award numbers R01GM126547 and R01GM127406) and from the US Army Research Office (award number W911NF-14-1-0411). The content is solely the responsibility of the authors and does not necessarily represent the official views of the NIH.

## Notes

### Competing Interest Statement

The authors have declared no competing interest.

